# Fast and accurate protein structure search with Foldseek

**DOI:** 10.1101/2022.02.07.479398

**Authors:** Michel van Kempen, Stephanie S. Kim, Charlotte Tumescheit, Milot Mirdita, Jeongjae Lee, Cameron L.M. Gilchrist, Johannes Söding, Martin Steinegger

## Abstract

As structure prediction methods are generating millions of publicly available protein structures, searching these databases is becoming a bottleneck. Foldseek aligns the structure of a query protein against a database by describing the amino acid backbone of proteins as sequences over a structural alphabet. Foldseek decreases computation times by four to five orders of magnitude with 86%, 88% and 133% of the sensitivities of DALI, TM-align and CE, respectively.

---

The recent developments in *in-silico* protein structure prediction at near-experimental quality [1, 2] are advancing structural biology and bioinformatics. The European Bioinformatics Institute already holds over 214 million structures predicted by AlphaFold2 [3] and the ESMAtlas contains over 617 million metagenomic structures predicted by ESMFold [4]. The scale of these databases poses challenges to state-of-theart analysis methods.

The most widely used approach to protein annotation and analysis is based on sequence similarity search [5–8]. The goal is to find homologous sequences from which properties of the query sequence can be inferred, such as molecular and cellular functions and structure. Despite the success of sequence-based homology inference, many proteins cannot be annotated because detecting distant evolutionary relationships from sequences alone remains challenging [9].

Detecting similarity between protein structures by 3D superposition offers higher sensitivity for identifying homologous proteins [10]. The availability of high-quality structures for any protein of interest allows us to use structure comparison to improve homology inference and structural, functional and evolutionary analyses. However, despite decades of effort to improve speed and sensitivity of structural aligners, current tools are much too slow to cope with today’s scale of structure databases.

Searching with a single query structure through a database with 100 M protein structures would take the popular TM-align [11] tool a month on one CPU core, and an all-versus-all comparison would take 10 millennia on a 1 000 core cluster. Sequence searching is four to five orders of magnitude faster: An all-versus-all comparison of 100 M sequences would take MMseqs2 [6] only around a week on the same cluster.

Structural alignment tools (reviewed in [12]) are slower for two reasons. First, whereas sequence search tools employ fast and sensitive prefilter algorithms to gain orders of magnitude in speed, no comparable prefilters exist for structure alignment. Second, structural similarity scores are non-local: changing the alignment in one part affects the similarity in all other parts. Most structural aligners, such as the popular TM-align, Dali, and CE [11, 13, 14], solve the alignment optimization problem by iterative or stochastic optimization. To increase speed, a crucial idea is to describe the amino acid backbone of proteins as sequences over a structural alphabet and compare structures using sequence alignments [15]. Structural alphabets thus reduce structure comparisons to much faster sequence alignments. Many ways to discretize the local amino acid backbone have been proposed [16]. Most, such as CLE, 3D-BLAST, and Protein Blocks, discretize the conformations of short stretches of usually 3 to 5 C_*α*_ atoms [17–19].

For Foldseek, we developed a type of structural alphabet that does not describe the backbone but rather tertiary interactions. The 20 states of the 3D-interactions (3Di) alphabet describe for each residue *i* the geometric conformation with its spatially closest residue *j*. 3Di has three key advantages over traditional backbone structural alphabets: (1) Weaker dependency between consecutive letters and (2) more evenly distributed state frequencies, both enhancing information density and reducing false positives (**Supplementary Table 1**). (3) The highest information density is encoded in conserved protein cores and the lowest in non-conserved coil/loop regions, whereas the opposite is true for backbone structural alphabets.

Foldseek (foldseek.com) (**Fig. 1a**) (1) discretizes the query structures into sequences over the 3Di alphabet, trained a 3Di substitution matrix (**Supplementary Table 2**), and then searches through the 3Di sequences of the target structures using the double-diagonal *k*-mer-based prefilter and gapless alignment prefilter modules from MMseqs2, our open-source sequence search software [6]. (2) High scoring hits are aligned locally using 3Di (default) or globally with TM-align (Foldseek-TM). The local alignment stage combines 3Di and amino acid substitution scores. The construction of the 3Di alphabet is summarized in **Fig. 1b** and **Supplemental Fig. 1-3**.

**FIG. 1.**
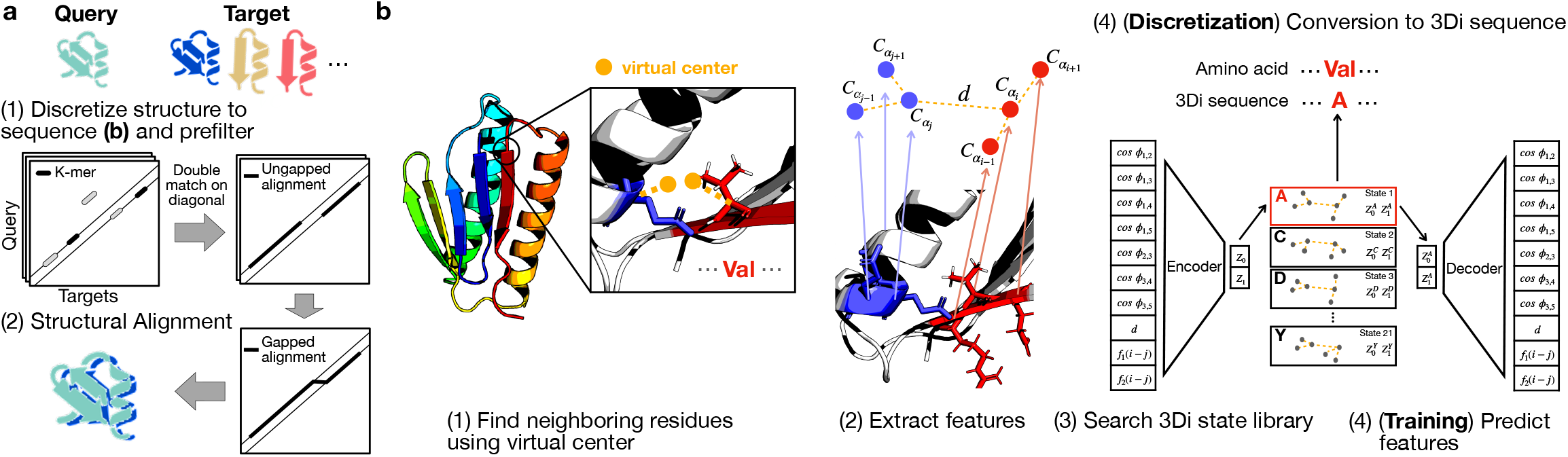
Foldseek workflow. (**a**) Foldseek searches a set of query structures through a set of target structures. (1) Query and target structures are discretized into 3Di sequences (see **b**). To detect candidate structures, we apply the fast and sensitive *k*-mer and ungapped alignment prefilter of MMseqs2 to the 3Di sequences, (2) followed by vectorized Smith-Waterman local alignment combining 3Di and amino acid substitution scores. Alternatively, a global alignment is computed with a 1.7 times accelerated TM-align version (see **Supplementary Fig. 4**). (**b**) Learning the 3Di alphabet: (1) 3Di states describe tertiary interaction between a residue *i* and its nearest neighbor *j*. Nearest neighbors have the closest virtual center distance (yellow). Virtual center positions (**Supplementary Fig. 1**) were optimized for maximum search sensitivity. (2) To describe the interaction geometry of residues *i* and *j*, we extract seven angles, the Euclidean C_*α*_ distance, and two sequence distance features from the six C_*α*_ coordinates of the two backbone fragments (blue, red). (3) These 10 features are used to define 20 3Di states by training a vector-quantized variational autoencoder [20] modified to learn states that are maximally evolutionary conserved. For structure searches, the encoder predicts the best-matching 3Di state for each residue.

To reduce high-scoring false positives and provide reliable E-values, we subtract the reversed query alignment score from the original score and apply a compositional bias correction within a local 40-residue sequence window (see “Pairwise local structural alignments”). E-values are calculated using an extreme-value score distribution, with parameters predicted by a neural network based on 3Di sequence composition and query length (see “E-Values”). Ranking of hits is determined by alignment bit score multiplied by the geometric mean of alignment TM-score and LDDT. Foldseek also reports the probability for each match to be homologous, based on a fit of true and false matches on SCOPe.

We measured the sensitivity and speed of Foldseek, six protein structure alignment tools, an alignment-free structure search tool (Geometricus [21]) and a sequence search tool (MMseqs2 [6]) on the SCOPe dataset of manually classified single-domain structures [22]. Clustering SCOPe 2.01 at 40 % sequence identity yielded 11 211 non-redundant protein sequences (SCOPe40). We performed an all-versus-all search and compared the tools’ performance for finding members of the same SCOPe family, superfamily, and fold (true positive matches, TPs) by measuring for each query the fraction of TPs out of all possible correct matches until the first false positive (FP), a match to a different fold (see “SCOPe Benchmark”).

We first measured the sensitivity to detect relationships at family and superfamily level by the area under the curve (AUC) of the cumulative ROC curve up to the first FP (**Fig. 2a, Supplementary Fig. 5**). Foldseek’s sensitivity is below Dali and TM-align, higher than the structural aligner CE, and much above the structural alphabet-based search tools 3D-BLAST and CLE-SW (**Fig. 2a**). In a precision-recall analysis, Foldseek-TM and Foldseek have the highest and third highest area under the precision-recall curve on each of the three levels (**Fig. 2b, Supplementary Fig. 5**). No-tably, Foldseek-TM improves over TM-align since its prefilter suppresses high-scoring false positives. Both sort hits by the average query and target length normalized TM-scores for best performance in the SCOPe benchmark.

**FIG. 2.**
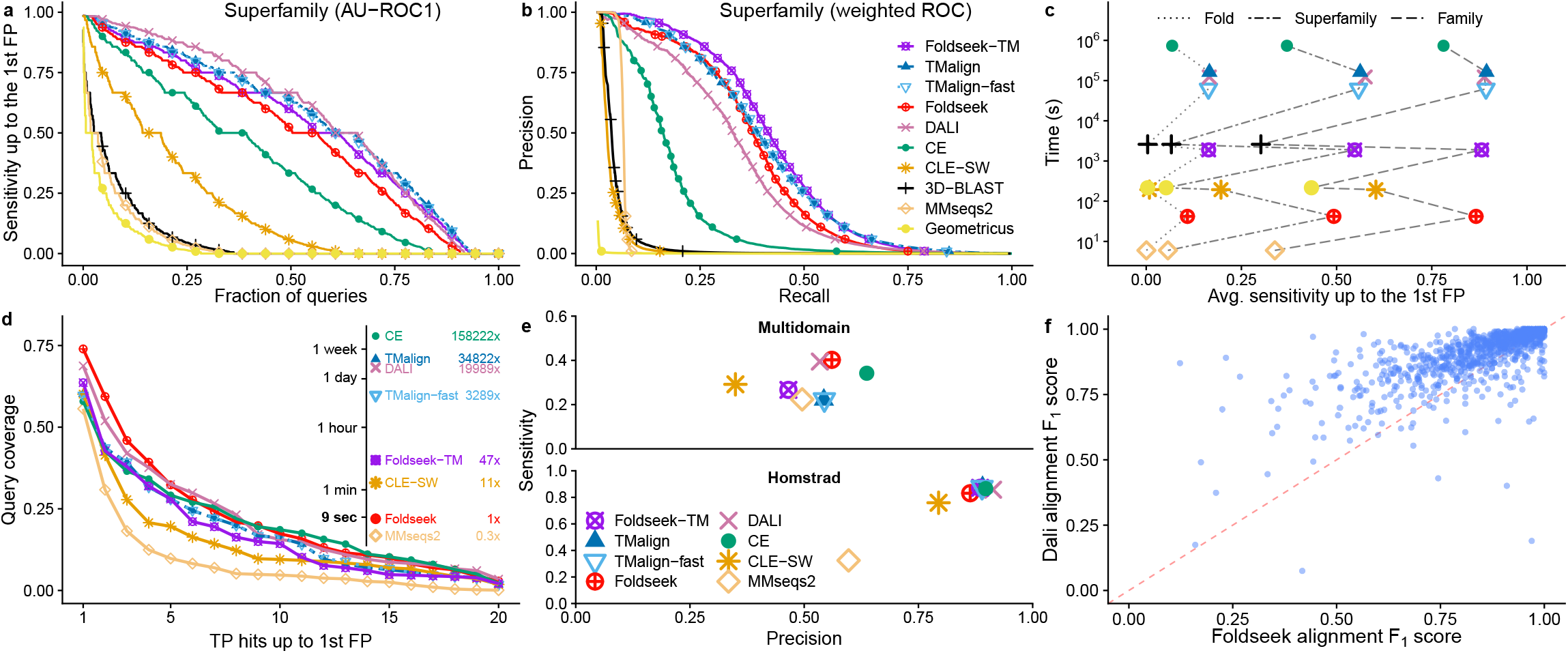
Foldseek reaches similar sensitivities as structural aligners at thousands of times their speed. (**a**) Cumulative distributions of sensitivity for homology detection on the SCOPe40 database of single-domain structures. True positives (TPs) are matches within the same superfamily, false positives (FPs) are matches between different folds. Sensitivity is the area under the ROC curve up to the first FP (see **Supplementary Fig. 5** for family and fold). (**b**) Precision-Recall curve of SCOPe40 superfamilies (see **Supplementary Fig. 5** for family and fold). (**c**) Avg. sensitivity up to the first FP for family, superfamily and fold versus total runtime on an AMD EPYC 7702P 64-core CPU for the all-versus-all searches of 11 211 structures of SCOPe40. (**d**) Search sensitivity on multi-domain, full-length AlphaFold2 protein models. 100 queries, randomly selected from AlphaFoldDB (v1), were searched against this database. Per-residue query coverage (y-axis) is the fraction of residues covered by at least *x* (x-axis) TP matches ranked before the first FP match. (**e**) Alignment quality for alignments of AlphaFoldDB (v1) protein models (top panel), averaged over the top five matches of each of the 100 queries. Sensitivity = TP residues in alignment / query length, precision = TP residues / alignment length. Reference-based alignment quality benchmark on HOMSTRAD alignments. (**f**) Alignment quality comparison between Foldseek and Dali for each HOMSTRAD family. The *F*_1_ score is the harmonic mean between sensitivity and precision.

Foldseek’s performance is comparable across all six secondary structure classes in SCOPe (**Supplementary Fig. 6**). On this small SCOPe40 benchmark set, Foldseek is more than 4,000 times faster than TM-align and Dali, and over 21,000 times faster than CE (**Fig. 2c**). On the much larger AlphaFoldDB (v1), where Foldseek approaches its full speed, it is around 184,600 and 23,000 faster than Dali and TM-align, respectively (see below).

We devised a reference-free benchmark to assess search sensitivity and alignment quality of structural aligners (see **Fig. 2d**) on a realistic set of full-length, multi-domain proteins. We clustered the AlphaFoldDB (v1) to 34,270 structures using BLAST and SPICi [23]. We randomly selected 100 query structures from this set and aligned them against the remaining structures. TP matches are those with a LDDT score [24] of at least 0.6 and FPs below 0.25, ignoring matches in-between. We set the LDDT thresholds according to the median inter- and intra-fold, -superfamily and -family LDDT scores of SCOPe40 alignments, see **Supplementary Fig. 7**. For other thresholds, see **Supplementary Fig. 8**. A domain-based sensitivity assessment would require a reference-based prediction of domains. To avoid it, we evaluated the sensitivity per residue. **Fig. 2d** shows the distribution of the fraction of query residues that were part of alignments with at least *x* TP targets with better scores than the first FP match. Again, Foldseek has similar sensitivity as Dali, CE, and TM-align and much higher than CLE-SW and MMseqs2.

We analyzed the quality of alignments produced by the top five matches per query. We computed the alignment sensitivity as the number of TP residues divided by the query length and the precision as the number of TP residues divided by the alignment length. TP residues are those with residue-specific LDDT score above 0.6, FP residues are below 0.25, residues with other scores are ignored. **Fig. 2e** shows the average sensitivity versus precision of the 100 ×5 structure alignments. Foldseek alignments are more accurate and sensitive than MMseqs2, CLE-SW, and TM-align, similarly accurate as Dali, and 13% less precise but 15% more sensitive than CE. In the reference-based HOMSTRAD alignment quality bench-mark [25], Foldseek performs slightly below CE, Dali, and TM-align (**Fig. 2e**). **Fig. 2f** shows the comparison between Foldseek and Dali in alignment quality for all HOMSTRAD families (see **Supplementary Fig. 9** for example alignments). To find potentially problematic high-scoring Foldseek FPs, we searched the set of unfragmented models in AlphaFoldDB (v1) with average predicted LDDT [1] ≥ 80 against itself. We inspected the 1,675 (of 133 813) highscoring FPs (score per aligned column ≥ 1.0, TM-score *<* 0.5), revealing queries with multiple structured segments but with incorrect relative orientations (**Supplementary Table 3, Supplementary Fig. 10**). The folded segments were correctly aligned by Foldseek. This illustrates that 3D aligners as TM-align may overlook homologous structures that are not globally superposable, whereas Foldseek (as well as the 2D aligner Dali) is independent of relative domain orientations and excels at detecting homologous multi-domain structures [12].

We developed a webserver (search.foldseek.com) for multi-database searches, including AlphaFoldDB (v1: Proteomes, Swiss-Prot), AlphaFoldDB (v4) at 50% sequence identity, ESMatlas-HQ, and PDB [26].

We compared Foldseek webserver, TM-align, and Dali using SARS-CoV-2 RdRp (PDB: 6M71_A [27]; 942 residues) in AlphaFoldDB (v1). Search times were 10 days for Dali, 33 h for TM-align, and 6 s for Foldseek, making it 180,000 and 23,000 times faster. All top 10 hits were known RdRp homologs (**Supplementary Table 4**).

The availability of high-quality structures for nearly every folded protein is transformative for biology and bioinformatics. Sequence-based analyses will soon be largely superseded by structure-based analyses. The main limitation in our view, the four orders of magnitude slower speed of structure comparisons, is removed by Foldseek.

## Acknowledgements

We thank Nicola Bordin, Ian Sillitoe and Christine Orengo for reporting issues and providing valuable feedback, Yang Zhang, Piotr Rotkiewicz and Marcin Wojdyr for making TM-align, PULCHRA and the Gemmi library freely accessible, and Do-Yoon Kim for creating the Foldseek logo.

M.S. acknowledges support from the National Research Foundation of Korea (NRF), grants [2019R1A6A1-A10073437, 2020M3A9G7103933, 2021R1C1C102065, 2021-M3A9I4021220], Samsung DS research fund and the Creative-Pioneering Researchers Program through Seoul National University. S.K. acknowledges support by NRF grant 2019R1-A6A1A10073437. J.S. acknowledges support by the German ministry for education and research (BMBF) (horizontal4meta). We used the compute cluster at the GWDG in Göttingen.

## Author contributions

M.K., S.K., J.S. & M.S. designed research. M.K., S.K., C.T., M.M. & M.S. developed code and performed analyses. M.K. and J.S. developed the 3Di alphabet. J.L. implemented the fast LDDT code, M.M. and C.L.M.G. developed the webserver. M.K., S.K., C.T., M.M., J.S. & M.S. wrote the manuscript.

## Competing interests

The authors declare no competing interests.

## METHODS

### Overview

Foldseek enables fast and sensitive comparison of large structure sets. It encodes structures as sequences over the 20-state 3Di alphabet and thereby reduces structural alignments to 3Di sequence alignments. The 3Di alphabet developed for Foldseek describes tertiary residue-residue interactions instead of backbone conformations and proved critical for reaching high sensitivities. Foldseek’s prefilter finds two *similar*, spaced 3Di *k*-mer matches in the same diagonal of the dynamic programming matrix. By not restricting itself to exact matches, the prefilter achieves high sensitivity while reducing the number of sequences for which full alignments are computed by several orders of magnitude. Further speed-ups are achieved by multi-threading and utilizing single instruction multiple data (SIMD) vector units. Owing to the SIMDe library (github.com/simd-everywhere/simde), Foldseek runs on a wide range of CPU architectures (x86_64, arm64, ppc64le) and operating systems (Linux, macOS). The core modules of Foldseek, which build on the MMseqs2 framework [28], are described in the following paragraphs.

### Create database

The `createdb` module converts a set of Protein Data Bank (PDB; [29]), macromolecular Crystallographic Information File (mmCIF) formatted files or Foldcomp compressed structure (FCZ, [30]) files into an internal Foldseek database format using the gemmi package (project-gemmi.github.io) or the Foldcomp library. The format is compatible with the MMseqs2 database format, which is optimized for parallel access. We store each chain as a separate entry in the database. The module follows the MMseqs2 `createdb` module logic. However, in addition to the amino acid sequence it computes the 3Di sequence from the 3D atom coordinates of the backbone atom and C_*β*_ coordinates (see “Descriptors for 3Di structural alphabet” and “Optimize nearest-neighbor selection”). Backbone atom and C_*β*_ coordinates are only needed for the nearest-neighbor selection. For C_*α*_-only structures, Foldseek reconstructs backbone atom coordinates using PULCHRA [31]. Missing C_*β*_ coordinates (e.g. in glycines) are defined such that the four groups attached to the C_*α*_ are arranged at the vertices of a regular tetrahedron. The 3Di and amino acid sequences and the C_*α*_ coordinates are stored in the Foldseek database. To save disk space, we optionally compress the C_*α*_ coordinates losslessly, beginning with three uncompressed 4-byte floating point C_*α*_ coordinates and storing all subsequent coordinates as two-bytes signed-integer differences [32]. If any difference is too large to be represented with a two-byte signed integer, we fall back to 4-byte floats for all C_*α*_ coordinates.

### Prefilter

The `prefilter` module detects double matches of similar spaced words (k-mers) that occur on the same diagonal. The k-mer size is dynamically set to *k* = 6 or *k* = 7 depending on the size of the target database. Similar k-mers are those with a 3Di substitution matrix score above a certain threshold, whereas MMseqs2 uses an amino acid substitution matrix to compute the similarity (see “3Di substitution score matrix”). The gapless double-match criterion suppresses hits to non-homologous structures effectively, as they are less likely to have consecutive k-mer matches on the same diagonal by chance. To avoid FP matches due to regions with biased 3Di sequence composition, a compositional bias correction is applied in a way analogous to MMseqs2 [33]. For each hit we perform an ungapped alignment over the diagonals with double, consecutive, similar k-mer matches and sort those by the maximum ungapped diagonal score. Alignments with a score of at least 15 bits are passed on to the next stage. We implemented an optional taxonomy-filter within the prefiltering step, to help users search through taxonomic subsets of the target database. After the gapless double-diagonal matching stage and before the ungapped alignment stage, we reject all potential target hits that do not lie within a taxonomic clade specified by the user.

### Pairwise local structural alignments

After the prefilter has removed the vast majority of non-homologous sequences, the `structurealign` module computes pairwise alignments for the remaining sequences using a SIMD accelerated Smith-Waterman algorithm [34, 35]. We extended this implementation to support amino acid and 3Di scoring, compositional bias correction, and 256-bit-wide vectorization. The score linearly combines amino acid and 3Di substitution scores with weights 1.4 and 2.1, respectively. We optimized these two weights and the ratio of gap-extend to gap open-penalty on ∼1 % of alignments (all-versus-all on 10% of randomly selected SCOPe40 domains). A compositional bias correction is applied to the amino acid and 3Di scores. To further suppress high-scoring FP matches, for each match we align the reversed query sequence against the target and subtract the reverse bitscore from the forward bitscore.

### Structural bit score

We rank hits by a “structural bit” score, that is the product of the bit score produce by the Smith-Waterman algorithm and the geometric mean of average alignment LDDT and the alignment TM-score.

### Fast alignment LDDT computation

To improve the LDDT score computation speed, we store the 3D coordinates of the query in a grid using spatial hashing. Each grid cell spans 15Å, which is the default radius considered for the LDDT computation. For each aligned query residue *i*, we compute the distances to all C_*α*_ atoms within a 15Å radius by searching all neighboring grid cells of the query residues grid cell. For each residue *j*, we compute the distance between the C_*α*_ atoms of *i* and *j* and the distance of the corresponding aligned target residues. Query and target distances for the aligned pairs are subtracted and the differences *d* are transformed into LDDT scores *s* = 0.25* ((*d <* 0.5) +(*d <* 1.0) +(*d <* 2.0) +(*d <* 4.0)). For each *i*, we obtain the means of the scores for all C*α* atoms *j* within the 15Å radius of *i*. The LDDT score is the mean of these means over all query residues *i*.

### E-Values

To estimate E-values for each match, we trained a neural network to predict the mean *μ* and scale parameter *λ* of the extreme value distribution for each query. The module `computemulambda` takes a query and database structures as input and aligns the query against a randomly shuffled version of the database sequences. For each query sequence the module produces *N* random alignments and fits to their scores an extreme-value (Gumbel) distribution. The maximum likelihood fitting is done using the Gumbel fitting function taken from HMMER3 (`hmmcalibrate`) [36]. To train the neural network, it is critical to use query and target proteins that include problematic regions such as structurally biased, disordered, or badly modeled regions that occur ubiquitously in full-length proteins or modeled structures. We therefore trained the network on 100 000 structures sampled from the AlphaFoldDB (v1). We trained a neural network to predict *μ* and *λ* from the amino acid composition of the query and its length (so a scrambled version of the query sequence would produce the same *μ* and *λ*). The network has 22 input nodes, 2 fully-connected layers with 32 nodes each (ReLU activation) and two linear output nodes. The optimizer ADAM with learning rate 0.001 was used for training. When testing the resulting E-values on searches with scrambled sequences, the log of the mean number of FPs per query turned out to have an accurately linear dependence on the log of the reported E-values, albeit with a slope of 0.32 instead of 1. We therefore correct the E-values from the neural network by taking them to the power of 0.32. We compared how well the mean number of FPs at a given E-value agreed with the E-values reported by Foldseek, MMseqs2, and 3D-Blast (**Supplementary Fig. 11**, see **Supplementary Fig. 12** for AlphaFoldDB). We considered a hit as FP if it was in a different fold and had a TM-score lower than 0.3. Furthermore, we ignored all cross-fold hits within the four- to eight-bladed *β*-propeller superfamilies (SCOPe b.66-b.70) and within the Rossman-like folds (c.2-c.5, c.27, c.28, c.30, and c.31) because of the extensive cross-fold homologies within these groups [37].

### Probability of true positive match

Foldseek computes for each match a simple estimate for the probability that the match is a true positive match given its structural bit score. Here, hits within the same superfamily are TP, hits to another fold are FP, and hits to the same family or to another superfamily are ignored. We estimate the structural bit score distributions of TP and FP hits (*p*(*score* |*TP*) and *p*(*score FP*)), which allow us to calculate the probability of a true positive 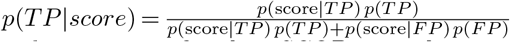. Both score distributions were fitted on SCOPe40 with a mixture model consisting of two gamma distributions (resulting in five parameters for each function). For the fitting, the function `gammamixEM` from the R package `mixtools` [38] was used. We excluded cross-fold hits between certain folds as in the E-value estimation. For example, Foldseek finds around the same number of FP and TP with a score of 51 in SCOPe40. The probability for a hit with score 51 is therefore 50%.

### Pairwise global structural alignments using TM-align

We also offer the option to use TM-align for pairwise structure alignment instead of the 3Di-based alignment. We implemented TM-align based on the C_*α*_ atom coordinates and made adjustments to improve the (1) speed and (2) memory usage. (1) TM-align performs multiple floating-point based Needleman-Wunsch (NW) alignment steps, while applying different scoring functions (e.g., score secondary structure, Euclidean distance of superposed structures or fragments, etc.) TM-align’s NW code did not take advantage of SIMD instructions, therefore, we replaced it by parasail’s [39] SIMD-based NW implementation and extended it to support the different scoring functions. We also replaced the TM-score computation using fast_protein_cluster’s SIMD based implementation [40]. Our NW implementation does not compute exactly the same alignment since we apply affine gap costs while TM-align does not **(Supplementary Fig. 4**). (2) TM-align requires 17 bytes × query length × target length of memory, we reduce the constant overhead from 17 to 4 bytes. If Foldseek is used in TM-align mode (parameter `--alignment-type 1`), TM-align is used for the alignment stage after the prefilter step, where we replace the reported E-value column with TM-scores normalized by the query length. The results are ordered in descending order by average TM-score by default.

### Descriptors for 3Di structural alphabet

The 3Di alphabet describes the tertiary contacts between residues and their nearest neighbors in 3D space. For each residue *i* the conformation of the local backbone around *i* together with the local backbone around its nearest neighbor *j* is approximated by 20 discrete states (see **Supplementary Fig. 3**). We chose the alphabet size *A* = 20 as a trade-off between encoding as much information as possible (large *A*, see **Supplementary Fig. 13**) and limiting the number of similar 3Di *k*-mers that we need to generate in the *k*-mer based prefilter, which scales with *A*^*k*^. The discrete single-letter states are formed from neighborhood descriptors containing ten features encoding the conformation of backbones around residues *i* and *j* represented by the C_*α*_ atoms (C_*α,i*−1_, C_*α,i*_, C_*α,i*+1_) and (C_*α,j*−1_, C_*α,j*_, C_*α,j*+1_). The descriptors use the five unit vectors along the following directions,

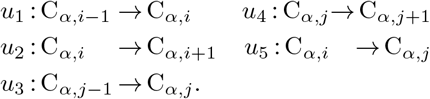

We define the angle between *u*_*k*_ and *u*_*l*_ as *ϕ*_*kl*_, so 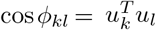. The seven features cos *ϕ*_12_, cos *ϕ*_34_, cos *ϕ*_15_, cos *ϕ*_35_, cos *ϕ*_14_, cos *ϕ*_23_, cos *ϕ*_13_, and the distance |C_*α,i*_ −C_*α,j*_| describe the conformation between the backbone fragments. In addition, we encode the sequence distance with the two features sign(*i*−*j*) min(|*i*−*j*|, 4) and sign(*i*−*j*) log(|*i*−*j*| + 1).

### Learning the 3Di states using a VQ-VAE

The ten-dimensional descriptors were discretized into an alphabet of 20 states using a variational autoencoder with vector-quantized latent variables (VQ-VAE) [41]. In contrast to standard clustering approaches such as k-means, VQ-VAE is a nonlinear approach that can optimize decision surfaces for each of its states. In contrast to the standard VQ-VAE, we trained the VQ-VAE not as a simple generative model but rather to learn states that are maximally conserved in evolution. To that end, we trained it with pairs of descriptors **x**_*n*_, **y**_*n*_ ∈ ℝ^10^ from structurally aligned residues, to predict the distribution of **y**_*n*_ from **x**_*n*_.

The VQ-VAE consists of an encoder and decoder network with the discrete latent 3Di state as a bottleneck in-between. The encoder network embeds the 10-dimensional descriptor **x**_*n*_ into a two-dimensional continuous latent space, where the embedding is then discretized by the nearest centroid, each centroid representing a 3Di state. Given the centroid, the decoder predicts the probability distribution of the descriptor **y**_*n*_ of the aligned residue. After training, only encoder and centroids are used to discretize descriptors. Encoder and decoder networks are both fully connected with two hidden layers of dimension 10, a batch normalization after each hidden layer and ReLU as activation functions. The encoder, centroids, and decoder have 242, 40, and 352 parameters, respectively. The output layer of the decoder consists of 20 units predicting *μ* and *σ*^2^ of the descriptors *x* of the aligned residue, such that the decoder predicts 𝒩 (*x* | *μ, Iσ*^2^) (with diagonal covariance).

We trained the VQ-VAE on the loss function defined in Equation (3) in [41] (with commitment loss = 0.25) using the deep-learning framework PyTorch (version 1.9.0), the ADAM optimizer, with a batch size of 512, and a learning rate of 10^−3^ over 4 epochs. Using Kerasify, we integrated the encoder network into Foldseek. The domains from SCOPe40 were split 80 %/20 % by fold into training and validation sets. For the training, we aligned the structures with TM-align, removed all alignments with a TM-score below 0.6, and removed all aligned residue pairs with a distance between their C_*α*_ atoms of more than 5 Å. We trained the VQ-VAE with 100 different initial parameters and chose the model that was performing best in the benchmark on the validation dataset (the highest sum of ratios between 3Di AUC and TM-align AUC for family, superfamily and fold level).

### 3Di substitution score matrix

We trained a BLOSUM-like substitution matrix for 3Di sequences from pairs of structurally aligned residues used for the “VAE-VQ training”. First, we determined the 3Di states of all residues. Next, the substitution frequencies between 3Di states were calculated by counting how often two 3Di states were structurally aligned. (Note that the substitution frequencies from state A to B and the opposite direction are equal.) Finally, the score 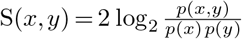 for substituting state x through state y is the log-ratio between the substitution frequency *p*(*x, y*) and the probability that the two states occur independently, scaled by the factor 2.

### 3Di alphabet cross-validation

We trained the 3Di alphabet (the VQ-VAE weights) and the substitution matrix by four-fold cross-validation on SCOPe40. We split the SCOPe40 dataset into four parts, such that all domains of each fold ended up in the same part of the four parts. 3Di alphabets were trained on three parts and tested on the remaining part, selecting each of the four parts in turn as a test set. The 80:20 split between training and validation sets to select the best alphabet out of the 100 VQ-VAE runs happens within the 3*/*4 of the cross-validation training data. **Supplementary Fig. 14** shows the mean sensitivity (black) and the standard deviation (gray area) in comparison to the final 3Di alphabet, for which we trained the 3Di alphabet on the entire SCOPe40 (red). No overfitting was observed, despite training 492 parameters (282 neural network, 210 substitution matrix entries). In **Fig. 2** we therefore show the benchmark results for the final 3Di alphabet, trained on the entire SCOPe40.

### Nearest-neighbor selection

To select nearest-neighbor residues that maximize the performance of the resulting 3Di alphabet in finding and aligning homologous structures, we introduced the virtual center *V* of a residue. The virtual center position is defined by the angle *θ* (*V* -C_*α*_-C_*β*_), the dihedral angle *τ* (*V* -C_*α*_-C_*β*_-N), and the length *l* (|*V* − C_*α*_|) (**Supplementary Fig. 1**). For each residue *i* we selected the residue *j* with the smallest distance between their virtual centers. The virtual center was optimized on the training and validation structure sets used for the VQ-VAE training by creating alphabets for positions with *θ* ∈ [0, 2*π*], *τ*∈ [ −*π, π*] in 45° steps, and *l* 1.53Å *k* : *k* 1, 1.5, 2, 2.5, 3 (1.53Å is the distance between C_*α*_ and C_*β*_). The virtual center defined by *θ* = 270°, *τ* = 0° and *l* = 2 performed best in the SCOPe benchmark.

This virtual center preferably selects long-range, tertiary interactions and only falls back to selecting interactions to *i* + 1 or *i* − 1 when no other residues are nearby. In that case, the interaction captures only the backbone conformation.

### SCOPe benchmark

We downloaded the SCOPe40 structures (available at wwwuser.gwdg.de/~compbiol/foldseek/scop40pdb.tar.gz).

The SCOPe benchmark set consists of single domains with an average length of 174 residues. In our benchmark, we compare the domains all-versus-all. Per domain, we measured the fraction of detected TPs up to the first FP. For family-, superfamily- and fold-level recognition, TPs were defined as same family, same superfamily and not same family, and same fold and not same superfamily, respectively. Hits from different folds are FPs.

### Evaluation SCOPe benchmark

After sorting the alignment result of each query (described in “Tools and options for benchmark comparison”), we calculated the sensitivity as the fraction of TPs in the sorted list up to the first FP, all excluding self-hits. For comparison, we took the mean sensitivity over all queries for family-, superfamily-, and fold-level classifications. We evaluated only SCOPe members with at least one other family, superfamily and fold member. We measure the sensitivity up to the 1th FP (ROC1) instead, for example, up to the 5th FP, because ROC1 actually better reflects the requirements for low false discovery rates in automatic searches.

Additionally, we plotted precision-recall curves for each tool (**Fig. 2b, Supplementary Fig. 5**). After sorting the alignment results by the structural similarity scores (as described in “Tools and options for benchmark comparison”) and excluding self-matches, we generated a weighted precision-recall curve for family-, superfamily-, and fold-level classifications (precision=TP/(TP+FP), recall=TP/(TP+FN)). All counts (TP, FP, FN) were weighted by the reciprocal of their family-, superfamily-, or fold size. In this way, folds, superfamilies, and families contribute linearly with their size instead of quadratically [37].

### Runtime evaluations on SCOPe and AlphaFoldDB

We measured the speed of structural aligners on a server with an AMD EPYC 7702P 64-core CPU and 1024 GB RAM memory. On SCOPe40, we measured or estimated the runtime for an all-versus-all comparison. To avoid excessive runtimes for TM-align, Dali, and CE, we estimated the runtime by randomly selecting 10 % of the 11 211 SCOPe domains as queries. We measured runtimes on AlphaFoldDB for searches with the same 100 randomly selected queries used for the sensitivity and alignment quality benchmarks (**Fig. 2d**,**e**). Tools with multi-threading support (MMseqs2 and Foldseek) were executed with 64 threads, tools without were parallelized by breaking the query set into 64 equally sized chunks and executing them in parallel.

### Reference-free multi-domain benchmarks

We devised two reference-free benchmarks that do not rely on any reference structural alignments. We clustered the AlphaFoldDB (v1) [42] using SPICi [43]. For this we first aligned all protein sequences all against all using an E-value threshold < 10^−3^ using BLAST (2.5.0+) [44]. SPICi produced 34,270 clusters from the search result. For each cluster we picked the longest protein as representative. We randomly selected 100 representatives as queries and searched the set of remaining structures. The top five alignments of all queries are listed at wwwuser.gwdg.de/~compbiol/foldseek/multi_domain_top5_alignments/.

For the evaluation, we needed to adjust the LDDT score function taken from AlphaFold2 [45]. LDDT calculates for each residue *i* in the query the fraction of residues in the 15 Å neighborhood which have a distance within 0.5,1,2,or 4 Å of the distance between the corresponding residues in the target [46]. The denominator of the fraction is the number of 15 Å-neighbors of *i* that are aligned to some residue in the target. This does not properly penalize non-compact models in which each residue has few neighbors within 15Å. We therefore use as denominator the *total* number of neighboring residues within 15 Å of *i*.

For the alignment quality benchmark (**Fig. 2e**), we classified each aligned residue pair as TP or FP depending on its *residue-wise* LDDT score, that is, the fraction of distances to its 15 Å neighbors that are within 0.5, 1, 2, and 4 Å of the distance to the corresponding residues in the query, averaged over the four distance thresholds. TP residues are those with a residue-wise LDDT score of at least 0.6 and FPs below 0.25, ignoring matches in-between. For the search sensitivity benchmark (**Fig. 2d**), TP residue-residue matches are those with an LDDT score of the query-target alignment of at least 0.6 and FPs below 0.25, ignoring matches in-between. (The LDDT score of the query-target alignment is the average of the residue-wise LDDT score over all aligned residue pairs.) The choice of thresholds is illustrated in **Supplementary Fig. 9**. The benchmark for other thresholds is shown in **Supplementary Fig. 10**.

### All-vs-all search of AlphaFoldDB with Foldseek

We downloaded the AlphaFoldDB (v1) [42] containing 365,198 protein models and searched it all-versus-all using Foldseek `-s 9.5 --max-seqs 2000`. For our second best hit analysis we consider only models with: (1) an average C_*α*_’s pLDDT greater than or equal to 80, and (2) models of non-fragmented domains. We also computed the structural similarity for each pair using TM-align (default options).

### Tools and options for benchmark comparison

Due to dataset overlap, we excluded methods from the benchmark that are likely to be overfitted on SCOPe. This applies to methods that trained many thousands of parameters (e.g. deep neural networks) with strong data leakage between training, validation, and test sets. For example several tools allowed up to 40% *sequence* identity between sets. The following command lines were used in the SCOPe as well as the multi-domain benchmark:

### Foldseek

We used Foldseek commit aeb5e during this analysis. Foldseek was run with the following parameters: `--threads 64 -s 9.5 -e 10 --max-seqs 2000`

### Foldseek-TM

For the Foldseek-TM benchmark we first run a regular 3Di/AA based Foldseek search using the following parameters: `--threads 64 -s 9.5 -e 10 --max-seqs 4000 --alignment-mode 1`. All hits passing are then aligned by Foldseeks’s `tmalign --tmalign-fast 1 --tmscore-threshold 0.0 -a`. We used Foldseek commit `aeb5e` during this analysis. We expose Foldseek-TM in our commandline interface as a search mode that combines regular Foldseek 3Di/AA based workflow with our TMalign implementation within the `tmalign` module.

### MMseqs2

We used the default MMseqs2 (release 13-45111) search algorithm to obtain the sequence-based alignment result. MMseqs2 sorts the results by e-value and score. We searched with: `--threads 64 -s 7.5 -e 10000 --max-seqs 2000`

### CLE-Smith-Waterman

We used PDB Tool v4.80 (github.com/realbigws/PDB_Tool) to convert the bench-mark structure set to CLE sequences. After the conversion, we used SSW [35] (commit ad452e) to align CLE sequences all-versus-all. We sorted the results by alignment score. The following parameters were used to run SSW: (1) protein alignment mode (`-p`), (2) gap open penalty of 100 (-o 100), (3) gap extend penalty of 10 (`-e` 10), (4) CLE’s optimized substitution matrix (`-a cle.shen.mat`), (5) returning alignment (`-c`). The gap open and extend values were inferred from DeepAlign [47]. The results are sorted by score in descending order.

`ssw_test -p -o 100 -e 10 -a cle.shen.mat -c`

### 3D-BLAST

We used 3D-BLAST (beta102) with BLAST+ (2.2.26) and SSW [35] (version ad452e). We first converted the PDB structures to a 3D-BLAST database using `3d-blast -sq_write` and `3d-blast -sq_append`. We searched the structural sequences against the database using `blastp` with the following parameters: (1) we used 3D-BLAST’s optimized substitution matrix (`-M 3DBLAST`), (2) number of hits and alignments shown of 12 000 (`-v 12000 -b 12000`), (3) E-value threshold of 1 000 (`-e 1000`) (4) disabling query sequence filter (`-F F`) (5) gap open of 8 (`-G 8`), and (6) gap extend of 2 (`-E 2`). 3D-BLAST’s results are sorted by E-value in ascending order:

`blastall -p blastp -M 3DBLAST -v 12000 -b 12000 -e 1000 -F F -G 8 -E 2`

For Smith-Waterman we used (1) gap open of 8 (2) gap extend of 2 and (3) returning alignments (`-c`) (4) using the 3D-BLAST’s optimized substitution matrix (-a 3DBLAST), (5) protein alignment mode (`-p`): `ssw_test -o 8 -e 2 -c -a 3DBLAST -p`. We noticed that the 3D-BLAST matrix with Smith-Waterman resulted in a similar performance to CLE: 0.717 0.230 0.011 for family-, superfamily- and fold-classification, respectively. We excluded 3D-BLAST’s measurement from the multi-domain benchmark since it produced occasionally high-scores (>10^7^) for single residue alignments.

### TM-align

We downloaded and compiled the `TMalign.cpp` source code (version 2019/08/22) from the Zhang group website. We ran the benchmark using default parameters and `-fast` for the fast version. TM-align reports two TM-scores: (1) normalized by the length of 1st chain (query) or (2) normalized by the length of the 2nd chain (target). We used the average of TM-scores normalized by the 1st chain (query) and 2nd chain (target) in all our analyses. We evaluated TMalign’s performance by sorting the results based on both the query TMscore and the minimum, maximum, and average TMscore for both the query and target. Our results showed that the average TMscore performed the best in our single domain benchmark.

Default: `TMalign query.pdb target.pdb`

Fast: `TMalign query.pdb target.pdb -fast`

### Dali

We installed the standalone DaliLite.v5. For the SCOPe40 benchmark set, input files were formatted in DAT files with Dali’s `import.pl`. The conversion to DAT format produced 11 137 valid structures out of the 11 211 initial structures for the SCOPe benchmark, and 34,252 structures out of 34,270 SPICi clusters. After formatting the input files, we calculated the protein alignment with Dali’s structural alignment algorithm. The results were sorted by Dali’s Z-score in descending order:

`import.pl –pdbfile query.pdb –pdbid PDBid –dat DAT dali.pl –cd1 queryDATid –db targetDB.list –TITLE systematic –dat1 DAT –dat2 DAT –outfmt “summary” –clean`

### CE

We used BioJava’s [48] (version 5.4.0) implementation of the combinatorial extension (CE) alignment algorithm. We modified one of the modules of BioJava under shape configuration to calculate the CE value. Our modified `CEalign.jar` file requires a list of query files, path to the target PDB files, and an output path as input parameters. This Java module runs an all-versus-all CE calculation, with unlimited gap size (`maxGapSize -1`) to improve alignment results [49]. The results were sorted by Z-score in descending order. For the multi-domain benchmark, we excluded 1 query that was running over 16 days. The Jar file of our implementation of CE calculation is provided (see “Code availability”).

`java -jar CEalign.jar querylist.txt`

`TargetPDBDirectory OutputDirectory`

### Geometricus

We included Geometricus [50] in the SCOPe benchmark as a representative of alignment-free tools, which are fast but can only find globally similar structures. Geometricus discretizes fixed-length backbone fragments (shape-mers) using their 3D moment invariants and represents structures as a fixed-length count vector over the shape-mers. To calculate the shape-mer-based structural similarity of the benchmark set, we used Caretta-shape’s Python implementation (1e3adb0) of multiple structure alignment (github.com/TurtleTools/caretta/caretta/multiple_alignment.py), which computes the BrayCurtis similarity between the Geometricus shape-mer vectors. Our modified version extracts structural information from the input files and generates all-versus-all pairwise structural similarity score as an output. We ran Geometricus on a single core because it would require substantial engineering efforts to implement parallelization on multiple cores. With an efficient multi-core implementation, Geometricus might be as fast as MMseqs2 on 64 cores. The Python code of our implementation of Geometricus is provided. `python runGeometricus_caretta.py -i querylist.txt -o OutputDirectory`

### HOMSTRAD alignment benchmark

The HOMSTRAD database contains expert-curated homologous structural alignments for 1032 protein families [51]. We downloaded the latest HOMSTRAD version (mizuguchilab.org/homstrad/data/homstrad_with_PDB_2022_Aug_1.tar.gz) and picked the pairwise alignments between the first and last members of each family, which resulted in structures of a median length of 182 residues. We used the same parameters as in the SCOPe and multi-domain benchmark. We forced Foldseek, MMseqs2, and CLE-Smith-Waterman to return an alignment by switching off the prefilter and E-value threshold. With the HOMSTRAD alignments as reference, we measured for each pairwise alignment the sensitivity (fraction of residue pairs of the HOMSTRAD alignment that were correctly aligned) and the precision (fraction of correctly aligned residue pairs in the predicted alignment). Dali, CE and CLE-Smith-Waterman failed to produce an alignment for 35, 1 and 1 out of 1032 pairs respectively, which were rated with a sensitivity of zero. The mean sensitivity and precision are shown in **Fig. 2e** and all individual alignments are listed in homstrad_alignments.txt at wwwuser.gwdg.de/~compbiol/foldseek/.

### Limitations of benchmarks

The SCOPe benchmark to measure search sensitivity only uses single-domain proteins as queries and targets (**Fig. 2a-c**). It therefore cannot assess the ability of tools to find local similarities, for example finding homologous domains shared between two multi-domain proteins. The alignment benchmark based on HOMSTRAD (**Fig. 2e**) has the same limitation, as the vast majority of proteins in HOMSTRAD have a single domain (median length = 182 residues). A drawback of our reference-free multi-domain benchmark is the need to choose thresholds for TPs and FPs (**Supplementary Fig. 7**).

### Prebuilt & ready-to-download databases

Foldseek includes the databases module to aid users with the download and set up of structural databases. Currently, we include the four variants of the AlphaFold Database (v4): UniProt (214M structures), UniProt50, a clustered database to 50% sequence identity and 90% bi-directional coverage using MMseqs2 (parameters `-c 0.9 --min-seq-id 0.5 --cluster-reassign`; 54M structures), Proteome (564k structures) and Swiss-Prot (542k structures). Additionally, we regularly build and offer a 100% sequence identity clustered PDB. The update pipeline is available in the `util/update_webserver_pdb` folder in the main Foldseek repository. These databases are hosted on Cloudflare R2 for fast downloading. We additionally link to and provide an automatic set-up procedure for the ESMatlas High-Quality Clu30 [52] database.

### Webserver

The Foldseek webserver is based on the MM-seqs2 webserver [53]. To allow for searches in seconds we implemented MMseqs2’s pre-computed database indexing capabilities in Foldseek. Using these, the search databases can be fully cached in system memory by the operating system and instantly accessed by each Fold-seek process, thus avoiding expensive accesses to slow disk drives. A similar mechanism was used to store and read the associated taxonomic information. The AlpaFoldDB/Uniprot50 (v4), AlphaFoldDB/Proteome (v4), AlphaFoldDB/Swiss-Prot (v4), ESMatlas High-Quality Clu30, and PDB100 require 191GB, 3.8GB, 3.4GB, 110GB, and 2.0GB RAM, respectively. The databases are kept in memory using `vmtouch` (github.com/hoytech/vmtouch). Databases are only required to remain resident in RAM, if Foldseek is used as a webserver. During batch searches, Foldseek adapts its memory use to the available RAM of the machine. We implemented a structural visualization using the NGL viewer [54] to aid the investigation of pairwise hits. Since we only store C_*α*_ traces of the database proteins, we use PULCHRA [31] to complete the backbone of these sequences, and also of the query if necessary, to enable a ribbon visualization [55] of the proteins. For a high quality superposition we use TM-align [56] to superpose the structures based on the Foldseek alignment. Both PULCHRA and TM-align are executed within the users’ browser using WebAssembly. They are available as `pulchra-wasm` and `tmalign-wasm` on the npm package repository as free open-source software.

### Structure prediction in the Webserver

We utilize the ESMatlas API to predict structures of user-supplied sequences that are at most 400 residues long. This enables sequence-to-structure searches in the webserver.

## Code availability

Foldseek is GPLv3-licensed free open source software. The source code and binaries for Foldseek can be downloaded at github.com/steineggerlab/foldseek. The webserver code is available at github.com/soedinglab/mmseqs2-app. The analysis scripts are available at: github.com/steineggerlab/foldseek-analysis.

## Data availability

Benchmark data is available at:

wwwuser.gwdg.de/~compbiol/foldseek

## Notes

https://foldseek.com

https://search.foldseek.com

### Competing Interest Statement

The authors have declared no competing interest.

### Summary of Updates

Foldseek has now integrated a scoring term that improves sensitivity by ∼30% in detecting super-families and folds. We additionally benchmarked Foldseek-TM. Additionally, include an alignment quality comparison benchmark with DALI.

