## Supplemental Material for "Fast and accurate protein structure search with Foldseek"

| Name | Mutual information [bit] |
|---|---|
| 3Di | 1.37 |
| CLE | 1.08 |
| 3D-BLAST | 1.07 |

**Supplementary Table 1: Mutual information per aligned residue pair** To estimate the information density of each structural alphabet, we measured the average mutual information between two structurally aligned residues $x$ and $y$. For this, we calculated the substitution frequencies for each alphabet as described in “3Di substitution score matrix”. Then, the mutual information equals $\sum_{x,y=1}^{20} p(x,y)\,\log_2 \frac{p(x,y)}{p(x)\,p(y)}$. We note that the mutual information is an upper limit on the mutual information per residue in sequences, since consecutive residues depend on each other, lowering the information content of sequences and thereby lowering the mutual information between structurally aligned sequences. Since 3Di sequences have weaker dependencies between consecutive residues than the CLE and 3D-BLAST alphabets (not shown), the gain in mutual information of 3Di over the two other alphabets is even higher than shown here.

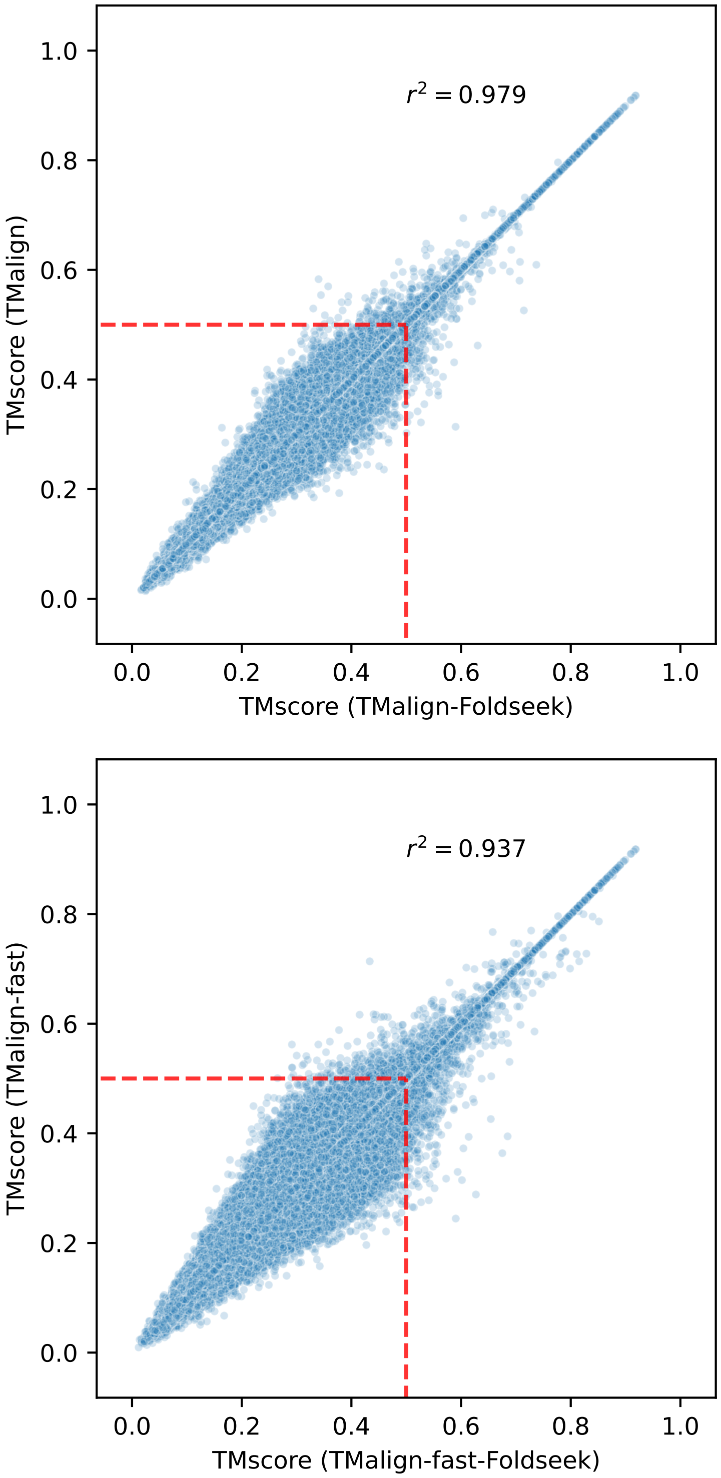


**Supplementary Figure** 1: **TMalign vs TMalign-Foldseek Benchmark** To compare our TMalign version (TMalign-Foldseek) with the original TMalign version, we sampled 1000 SCOPe domains and performed an all-vs-all alignment in both TMalign modes (normal and fast). TMalign-Foldseek was 1.7 times faster (253 minutes) compared to TMalign (438 minutes). When using the fast mode, this factor remained the same between TMalign-Foldseek (75 minutes) and TMalign (131 minutes). The two figures show the relation between the TMscores of the two implementations in both normal and fast mode respectively. Note that TMscores below 0.5 (red dashed line) indicate that the structures are not from the same fold.

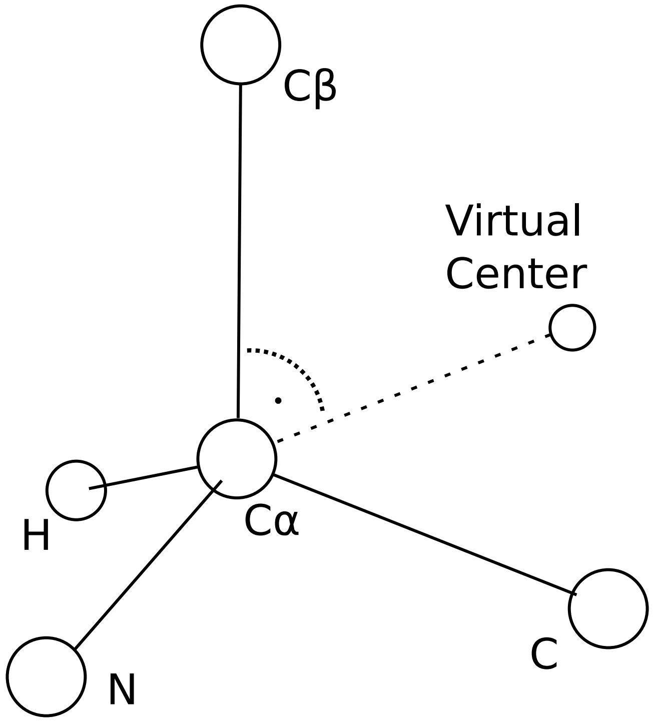


**Supplementary Figure** 2: **3Di virtual center.** During the transformation of structures into 3Di sequences, the virtual centers of residues are used to determine interacting residues. The optimized virtual center lies on the plane defined by the atoms N, $C_\alpha$, and $C_\beta$. Moreover, $C_\beta$, $C_\alpha$, and the virtual center form an angle of 90°. The distance between the virtual center and $C_\alpha$ equals twice the distance between $C_\beta$ and $C_\alpha$. For glycines, the $C_\beta$ is approximated by assuming that the $C_\beta$, $C_{backbone}$, and N atoms are arranged at the vertices of a regular tetrahedron with $C_\alpha$ at its centroid, and a centroid to vertex distance of 1.5336 Å.

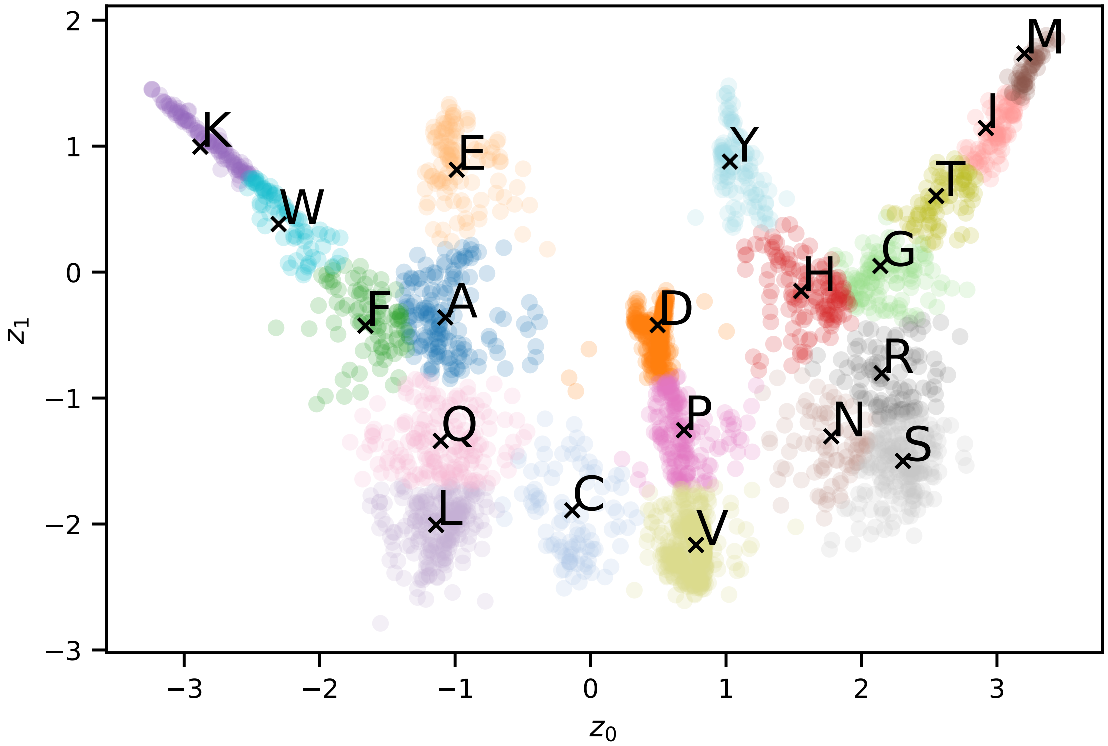


**Supplementary Figure** 3: **Latent space representation learned by encoder network** The encoder network of the VQ-VAE encodes the 3Di descriptor of a residue into a two-dimensional representation. Here, we show this latent space representation of 3000 sampled residues. Each circle represents a residue and is colored according to its nearest centroid (x), which discretizes the residue to a 3Di state.

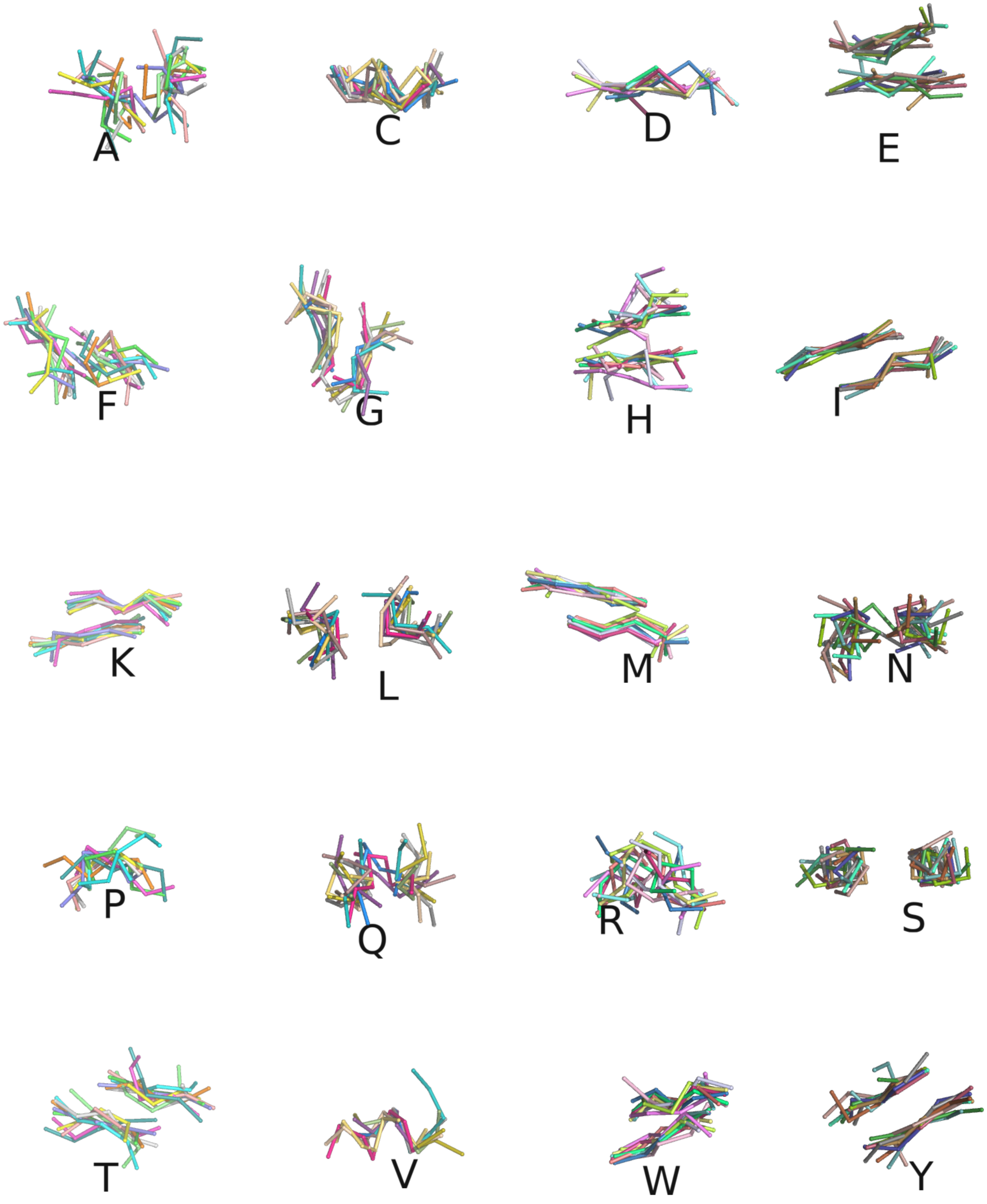


**Supplementary Figure** 4: **3Di state visualizations** Each 3Di state represents a conformation between two three-residue backbone fragments. To visualize this conformation, we sampled and aligned ten fragment pairs for each state, where the paired fragments have the same color. Here, five-residue fragments are shown, however the 3Di states describes only the conformation of the inner three-residue fragments.

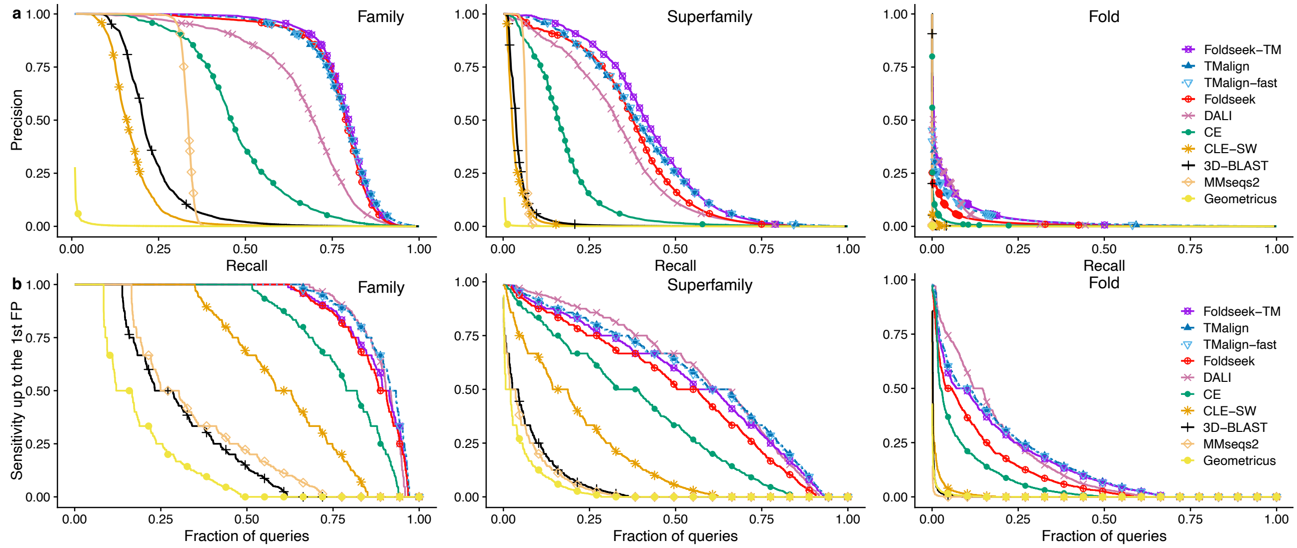


**Supplementary Figure 5: Precision-Recall curves and cumulative distributions of sensitivity for the SCOPe benchmark** We measured the sensitivity of Foldseek and seven structure alignment tools on the SCOPe40 dataset by plotting precision-recall curves and the cumulative distributions of sensitivity up to the first FP on family, superfamily, and fold level.

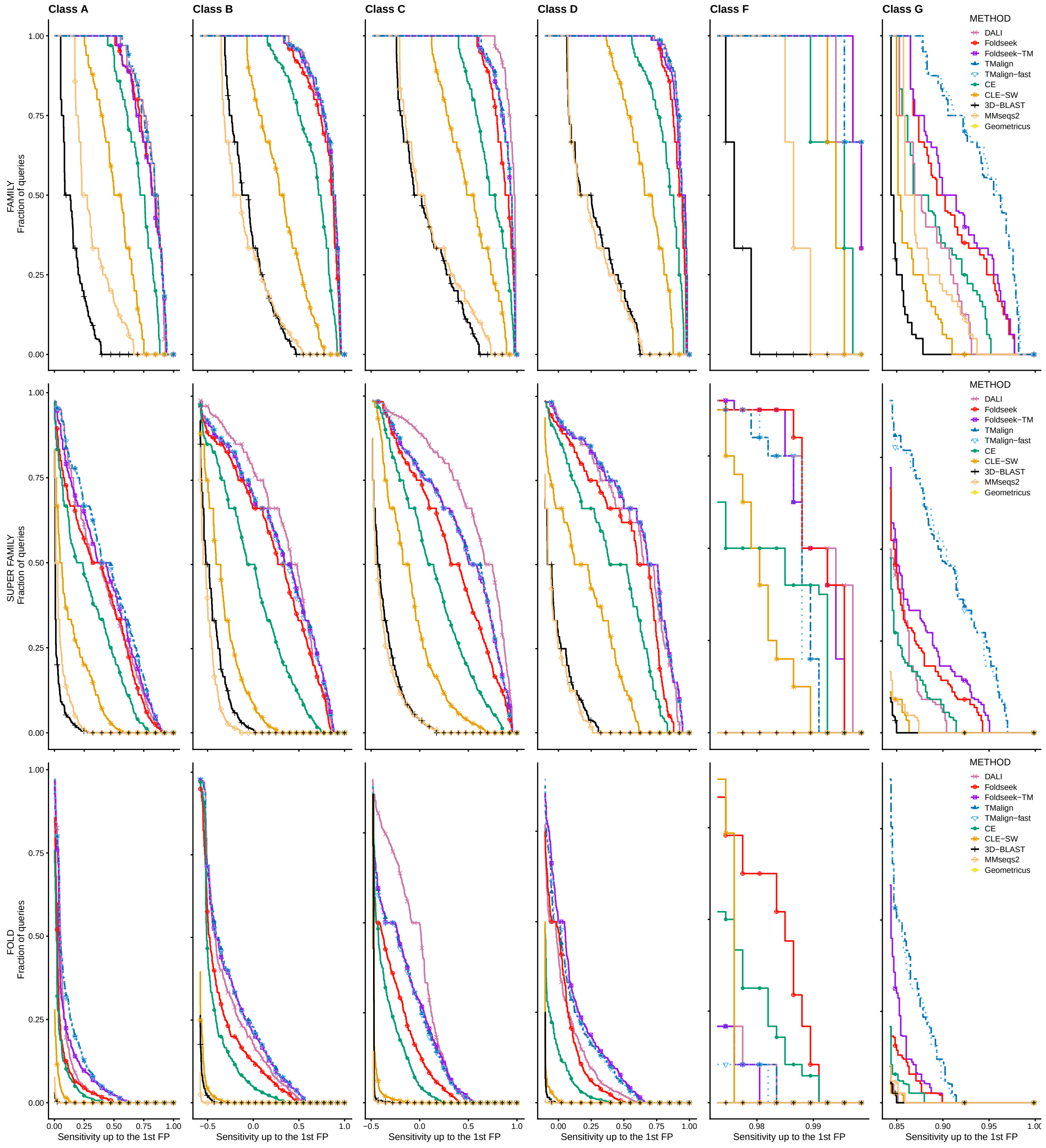


**Supplementary Figure 6**: **SCOPe benchmark by class.** We measured the sensitivity of Foldseek and six structure alignment tools on the SCOPe40 dataset, which are divided into six SCOPe classes: (1) Class A contains all alpha proteins, (2) class B has all beta proteins, (3) class C consists alpha-beta proteins with mainly parallel beta strands (a/b), (4) class D alpha-beta proteins with mainly antiparallel beta strands (a+b), (5) class F consists of membrane and cell surface proteins and peptides, and (6) class G contains small proteins.

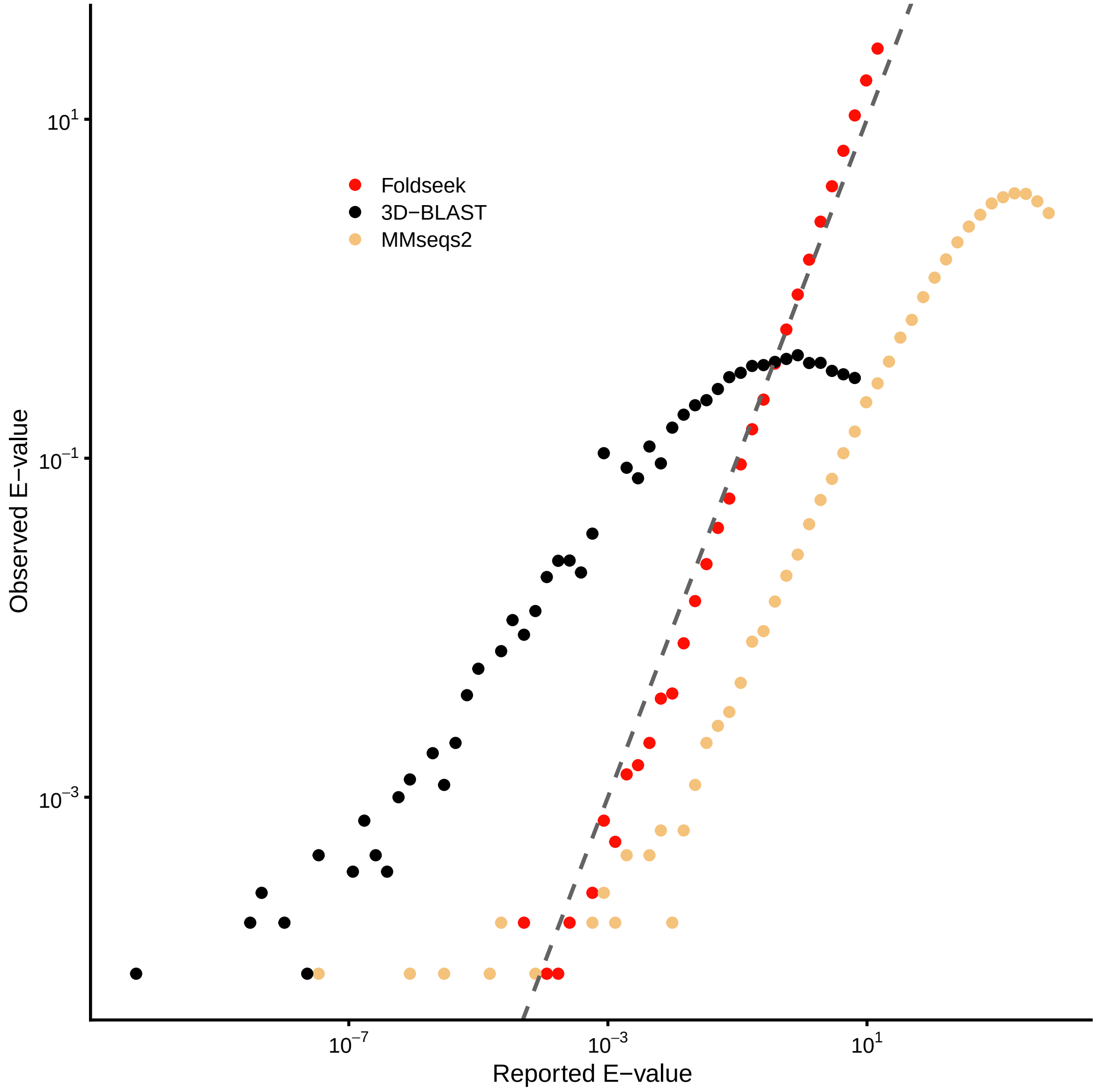


**Supplementary Figure 7**: **Accuracy of reported E-values** Mean number of FPs per query below the reported E-value threshold measured on SCOP (see **Supplementary Fig. 8** for AlphaFold DB structures)

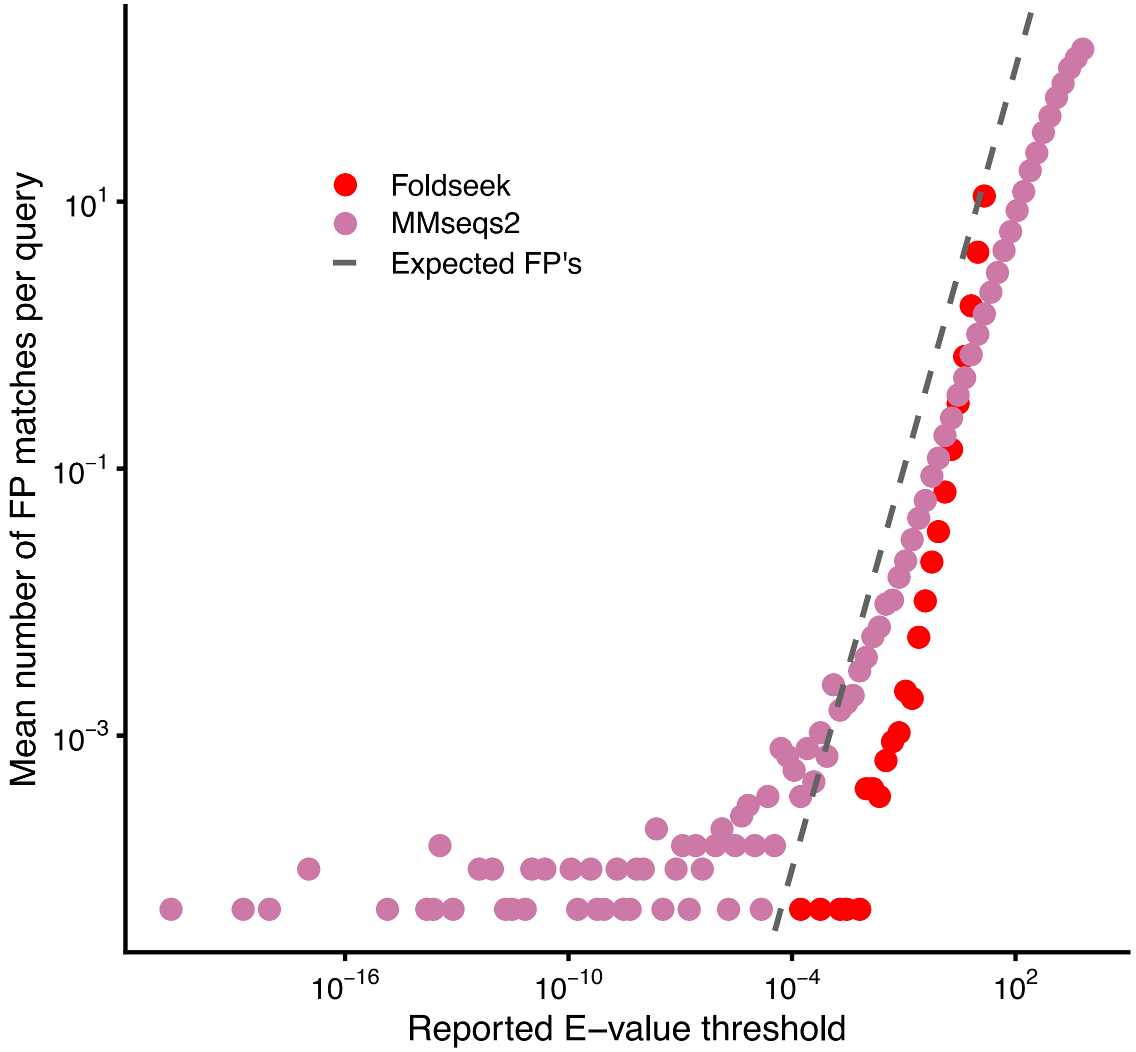


**Supplementary Figure** 8: **E-values for shuffled sequences from AlphaFoldDB** As described in the Methods section, we trained a neural network to predict the Gumbel distribution parameters from the input sequence composition. Here, we show the mean number of FP matches per query versus the reported E-value threshold for 100 000 randomly sampled sequences from the AlphaFoldDB and shuffled those as described earlier. E-values are shown for Foldseek and MMseqs2.

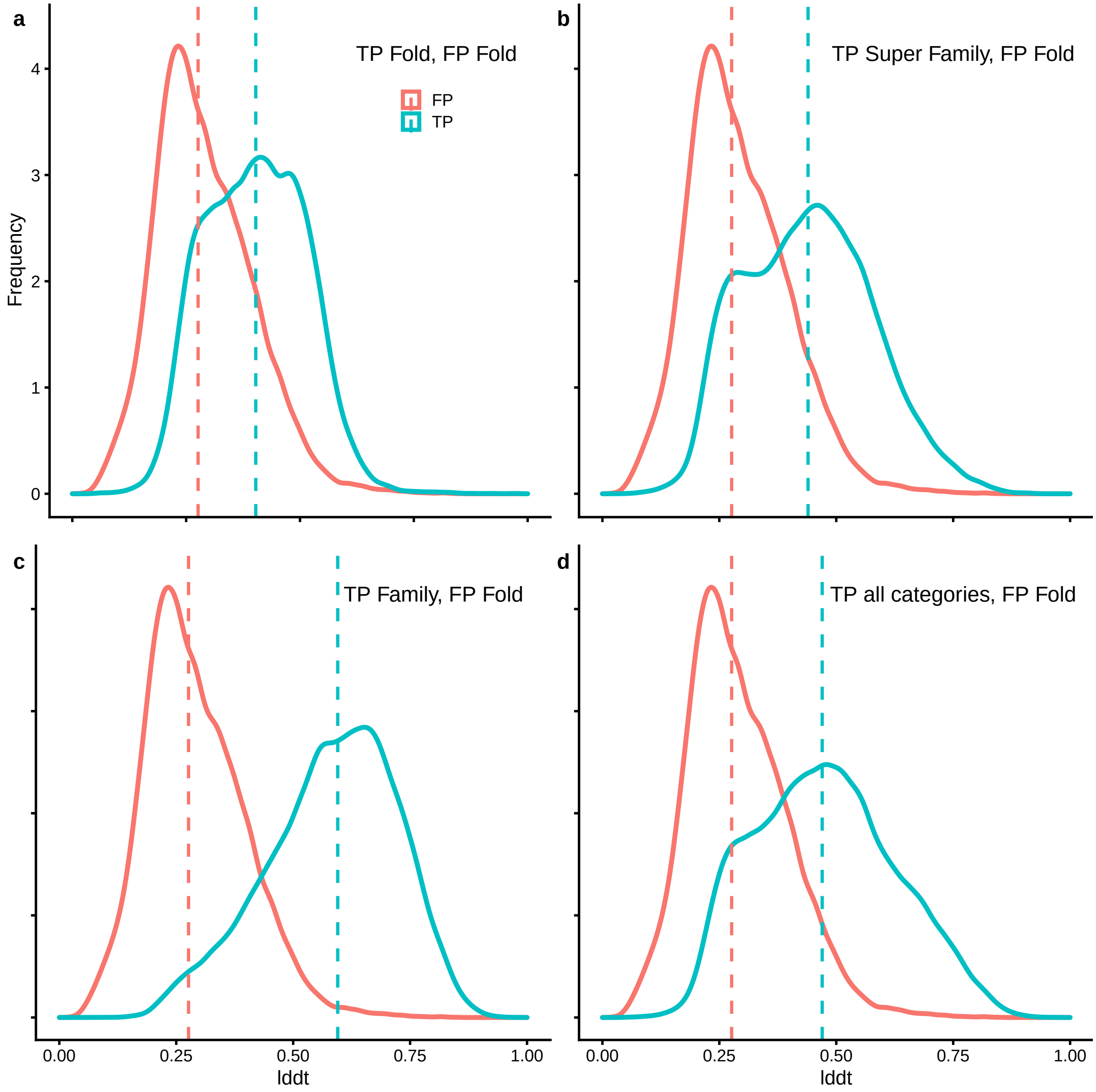


**Supplementary Figure** 9: **SCOPe40 TP and FP average LDDT per alignment distribution.** Average LDDT distribution of 10,000 randomly chosen pairwise alignments of TP and FP hits for TM-align, Foldseek, Dali, and CLE. FP shown for folds (a-d), TP shown for folds (a), superfamilies (b), families (c), and all categories together (d).

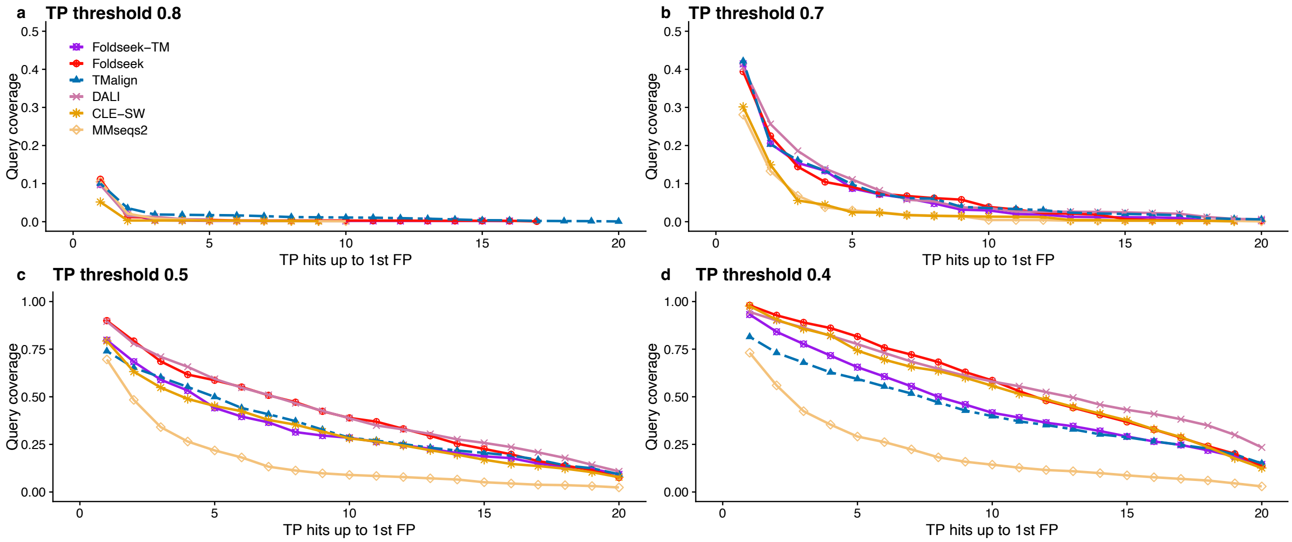


**Supplementary Figure** 10: **AlphaFold coverage benchmark at different TP thresholds.** This figure shows the same benchmark as in **Fig 2d**, but with different LDDT thresholds for TP hits, while the FP threshold remains at 0.25.

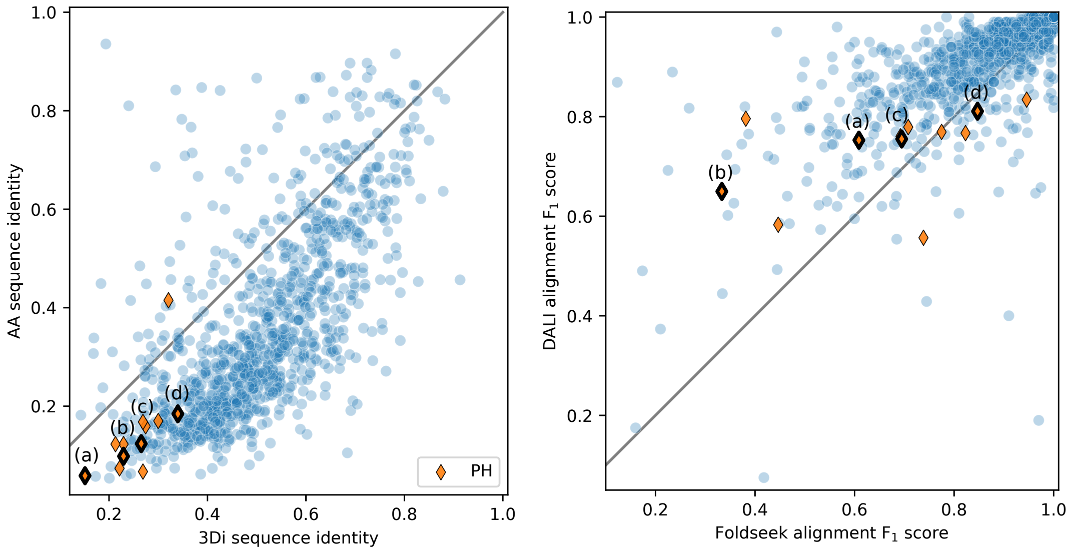


(1) **Left**: Comparison of sequence identities of amino acid (AA) and 3Di sequences in HOMSTRAD reference alignments. For each HOMSTRAD family, the alignment between the first and last member was used (blue circles). We selected the HOMSTRAD family `PH` as its members cover a large range in both amino acid sequence identity and Foldseek's alignment quality. The orange diamonds represent alignments within this family (between `1dro` and all other family members). Next, we selected a series of alignments with increasing amino acid sequence identity ((a) to (d)). For each alignment, the HOMSTRAD reference alignment and Foldseek's alignment are shown below. **Right**: Alignment quality comparison on HOMSTRAD families (blue circles, also shown in **Fig. 2f**) between Foldseek and Dali, which performed best in the HOMSTRAD benchmark. For each family, the alignment quality between the first and last family member is described by the $F_1$ score, which is the harmonic mean of sensitivity and precision. The average sensitivity and precision for all methods is shown in **Fig. 2e**. 46 of 1032 families are not shown, because Dali did not return an alignment.

(a) 1dro - 1mai (sensitivity = 0.57, precision = 0.66, $F_1$ score = 0.61)

HOMSTRAD alignment: (RMSD=5.4Å for 99 residues, seq.id. = 6% (AA), 15% (3Di))

```
1dro        1 D------PDDDDDAAAWDQFWKQKFDDDDDAVPHPDHIDGATWDDAAA---KTFGARDCAVVVVDVQDHPVVTDMGGALQ
              |               |+++|++ +          +++++ +|+       |+|   |         |++ |  +++  +
1mai        1 DLVPDPLNVVQQV---WAFWWWFDAQ---------PRTHTKIWHQHP-VNFKIFIHDDDP-------DDDRVVRIDTLVF

1dro       72 W-AWDADC------DDDQSNL---KIWTGGPP-RIIIITRDDDNVVRVVSVVGRHCRNN
              + |+++++      +         ++||+|++   |++++++|+| ++   | + ++|
1mai       61 FDAKAFAQPDPRCVHRVPVADRLFWMWTDGDDPDDITTIGHPDSVNSCSVRVVSVVSND
```

Foldseek alignment: (RMSD=5.5Å for 85 residues)

```
1dro       11 WDQFWKQKFDDDDDAVPHPDHIDGATWDDAA--AKTFGARDCAVVVVDVQDHPVVTDMGGALQW-AWDADCDD-------
              |+++|++      || |   ++++ +|+     +|+|   |      |  |+ ||  +++  ++ |++++++|
1mai       14 WAFWWWF------DAQP---RTHTKIWHQHPVNFKIFIHDD------DPDDDRVVR-IDTLVFFDAKAFAQPDPRCVHRV

1dro       81 --DQSNLKIWTGG--PPRIIIITRDDDNVVRVVSVV
                + +  ++||+|  |+ |++|+ ++|+| ++   |
1mai       78 PVADRLFWMWTDGDDPDDITTIG-HPDSVNSCSVRV
```

(2)

(b) 1dro - 1b55A (sensitivity = 0.32, precision = 0.35, $F_1$ score = 0.33)

HOMSTRAD alignment: (RMSD=5.6Å for 100 residues, seq.id. = 10% (AA), 23% (3Di))

```
1dro       6 DDAAAWDQFWKQKFDDDDDAVPHPDHIDGATWDDAAA-KTFGARD-CAVVVVDVQDHPVVTDMGGALQW-AWDADC----
             ||  +++ +|+    +|+    ++ +||+ +| |+   ++++|++     | |  + +   |+++ +|| ++| +
1b55a      1 DDWPDKAWWWWWDPCPDP---VDDTDIDTWIWTDDL-FWIWTAHADPV--VGDGHHTD---DIDTLQQWQDKDWAPFDDD

1dro      79 DDDQSNL--KIWTGGPP--------------------RIIIITRDDDNVVRVVSVVGRHCRNNDD---------------
             || |                                   |||+ + | || ||  | + ++  ++
1b55a     72 DDQQQDDDD--------DPDVVQQRRRFNTWMWTAGDVAIIIITGSDPVVVVVVRVVSCVSHVPHDRHDQAAARYDQPDQ

1dro     122 -------------------D
                                |
1b55a    144 GGPVVRDRRRHDGHPDGDDD
```

Foldseek alignment: (RMSD=6.7Å for 92 residues)

```
1dro      14 FWKQKFDDDDDAVPHPDHIDGATWDDAAAKTFGARDCAVVVVDVQDHPVVT-DMGGALQW-AWD---ADCDDDQS-----
             +|+ +  |+ ++  ++ +||+ +| |+ +++++|+  |  ||+  +|   | |+++ +|| ++|   +| ||||
1b55a      8 WWWWW--DPCPDPVDDTDIDTWIWTDDLFWIWTAH--ADPVVGDGHH---TDDIDTLQQWQDKDWAPFDDDDDQQQDDDD

1dro      84 -----------NLKIWTGGPPRIIIITRDDDNVV---RVVSVV
                        | ++|| |+  |||||+ |+ ||   ||||+|
1b55a     81 DPDVVQQRRRFNTWMWTAGDVAIIIITGSDPVVVVVVRVVSCV
```

(3)

(c) 1dro - 1dyna (sensitivity = 0.65, precision = 0.74, $F_1$ score = 0.69)

HOMSTRAD alignment: (RMSD=4.3Å for 101 residues, seq.id. = 12% (AA), 27% (3Di))

```
1dro       6 DDAAAWDQFWKQKFDDDDDAVPHPDHIDGATWDDAAA-KTFGARDCAVVVVDVQDHPVVTDMGGALQWAWDADC-DDDQS
             ||  +++ +|+|  |        |  |+|  +| |+   ++++|+|+        +  |   |+ +  |+|+ ++  ||+
1dyna      1 DDWLDKAKWWFQQDD------VVPVHTDIWIWIDDL-FWIWTANDVVRP-----GIP---DIFTLAQKAKEFPPCDDPDQ

1dro      84 NLKIWTGGPP-------RIIIITRDDDNVVRVVSVVGRHCRNNDD
               |+|+ + |       + | |+ | | || ||  | + ++
1dyna     66 WGKMWIARNPDQARPDNHSIGIITDNDPVVVVVVQVSSVVSPHYY
```

Foldseek alignment: (RMSD=4.2Å for 89 residues)

```
1dro      10 AWDQFWKQKFDDDDDAVPHPDHIDGATWDDAAAKTFGARDCAVVVVDVQDHPVVT-DMGGALQWAW-DADCDDDQSNLKI
             +++ +|+|      |+|    |+|  +| |+ +++++|+|+      | ++   + |+ +  |+|+   +|||+    |+
1dyna      5 DKAKWWFQ-----QDDVVPV-HTDIWIWIDDLFWIWTANDV------VRPG---IPDIFTLAQKAKEFPPCDDPDQWGKM

1dro      88 WTGGPP--------RIIIITRDDDNVVRVVSVVGR
             |+ + |        +| ||| | | || ||  | +
1dyna     70 WIARNPDQARPDNHSIGIIT-DNDPVVVVVVQVSS
```

(4)

(d) 1dro - 1qqgA (sensitivity = 0.77, precision = 0.94, $F_1$ score = 0.85)

HOMSTRAD alignment: (RMSD=3.4Å for 94 residues, seq.id. = 18% (AA), 34% (3Di))

```
1dro        7 DAAAWDQFWKQKFDDDDDAVPHPDHIDGATWDDAAA-------KTFGARDCAVVVVDVQDHPVVTDMGGALQW-AWDADC
              | ++|| +  |  +             + ++  +||       ++| ++|  +|||   | |   |+|+  |+ +||+|+
1qqga       1 DWDDWDWWAQQ--P----------VRQTWIKTFDAADPPPHATWIFIDNDDVCVVVVPDDTP---DIGHLVQFPDWDFDQ

1dro       79 DDDQSNLKIWTGGPP-RIIIITRDDDNVVRVVSVVGRHCRN
              |+   + ||||+||    | |++++| ||+||  | + +|
1qqga      66 DPR-ARTKIWTHGP-P-DIGIIGHPDVVVSVVVSVSSVVRD
```

Foldseek alignment: (RMSD=3.2Å for 76 residues)

```
1dro       34 GATWDDAAA-------KTFGARDCAVVVVDVQDHPVVTDMGGALQW-AWDADCDDDQSNLKIWTGGPPRIIIITRDDDNV
              + ++  +||       ++| ++|  +|||   | |   |+|+  |+ +||+|+|+ + + ||||+||| |  |++++| |
1qqga      16 TWIKTFDAADPPPHATWIFIDNDDVCVVVVPDDTP---DIGHLVQFPDWDFDQDPRA-RTKIWTHGPPDI-GIIGHPDVV

1dro      106 VRVVSVVGR
              |+||  | +
1qqga      91 VSVVVSVSS
```

(5)

**Supplementary Figure** 11: **Foldseek alignment quality of example alignments** Subfigure (1) shows the sequence identities and alignment qualities of the four selected alignments in context of all HOMSTRAD alignments. The alignments are shown in subfigures (2) to (5). For each example alignment, the 3Di sequences aligned by HOMSTRAD and by Foldseek are shown. The sequence identities of aminco acid (AA) and 3Di sequence in the HOMSTRAD alignment are provided. The root mean square deviation (RMSD) was calculated for both alignments. The symbol | denotes an alignment of identical 3Di states, and the symbol + denotes an alignment between 3Di states with positive substitution score.

| | Query | Target | Foldseek | | TM align |
|---|---|---|---|---|---|
| | | | Bit per column | E-value | TM score |
| **Model** | AF-P0AEC3-F1 | AF-P30855-F1 | 3.021 | 3.459E-43 | 0.291 |
| **Protein Function** | Aerobic respiration control sensor protein ArcB | Sensor protein EvgS | | | |
| | EC: 2.7.13.3 | EC: 2.7.13.3 | | | |
| **Structure** | 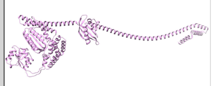 | 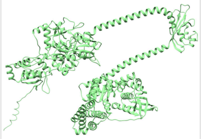 | | | |
| **Sequence Length** | 778 | 1197 | | | |
| **Avg. pLDDT** | 81.193 | 82.329 | | | |
| **Predicted Aligned Error** | 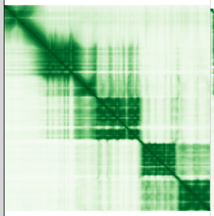 | 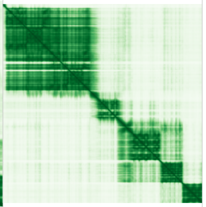 | | | |
| **Model** | AF-Q4CPQ6-F1 | AF-Q4CLF8-F1 | 5.856 | 4.293E-61 | 0.233 |
| **Protein Function** | Calpain cysteine peptidase, putative | Calpain-like cysteine peptidase, putative | | | |
| | - | - | | | |
| **Structure** | 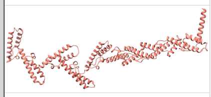 | 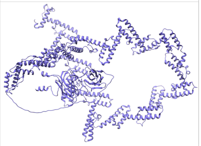 | | | |
| **Sequence Length** | 1753 | 631 | | | |
| **Avg. pLDDT** | 82.150 | 89.192 | | | |
| **Predicted Aligned Error** | 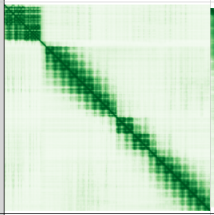 | 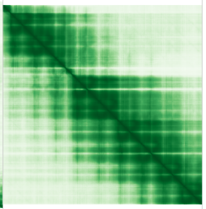 | | | |
| **Model** | AF-Q54P41-F1 | AF-Q54QK8-F1 | 1.938 | 1.146E-38 | 0.367 |
| **Protein Function** | EGF-like domain-containing protein | EGF-like domain-containing protein | | | |
| | - | - | | | |
| **Structure** | 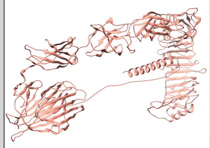 | 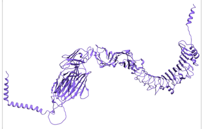 | | | |
| **Sequence Length** | 992 | 1064 | | | |
| **Avg. pLDDT** | 80.493 | 86.856 | | | |
| **Predicted Aligned Error** | 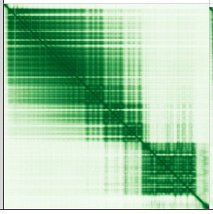 | 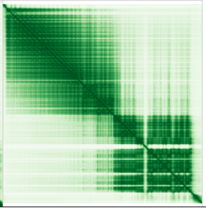 | | | |

**Supplementary Table** 2: **Outliers in Foldseek's top hits in AlphaFoldDB** Three hand-picked outliers from the scatter plot in **Supplemental Fig. 12** are shown here. The outliers were selected from examples where Foldseek has a hit with a score greater 1.0 bits per column and an TM-align score below 0.5. These three models taken from the AlphaFoldDB contain well predicted local structures (with high average pLDDTs of over 80). However, the global structure is not well predicted (visible in the light green areas of the Predicted Aligned Error). This shows Foldseek's ability to detect remote homologies that are difficult to find by TM-align's global alignment.

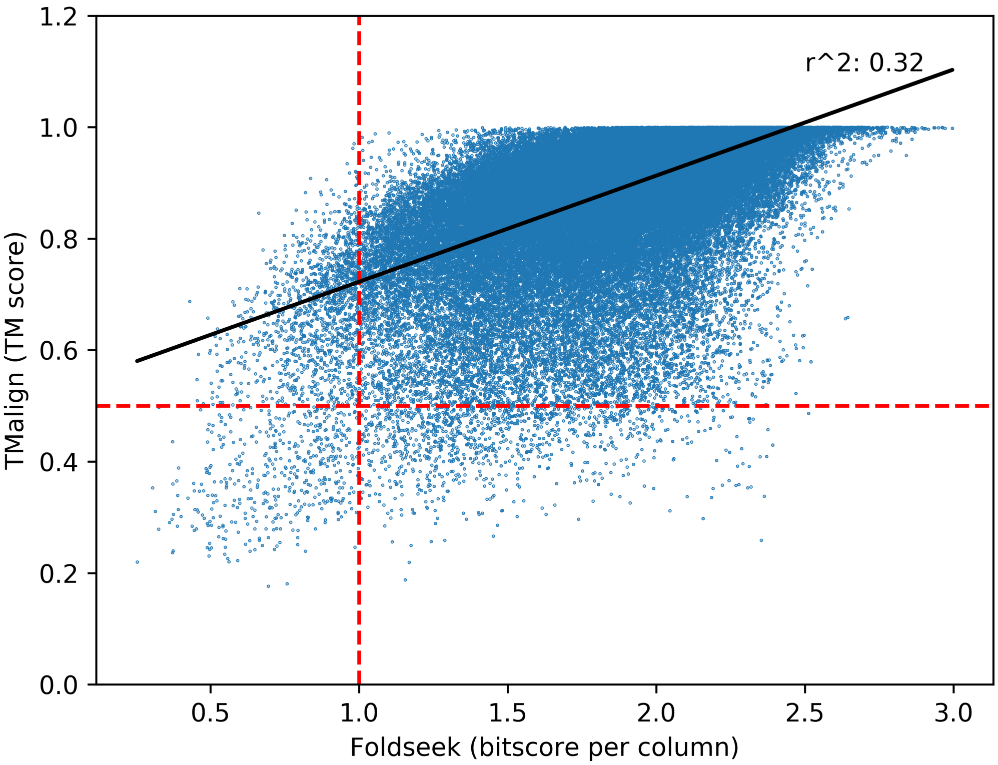


**Supplementary Figure** 12: **Structural similarity of Foldseek's top hits in AlphaFoldDB** After running an all-versus-all alignment on AlphaFoldDB with Foldseek, the top hit (sorted by Foldseek's E-value) for each query was collected and structural similarity was calculated by TM-score as a comparison. The thresholds for each tool (Foldseek: $>$ 1.0 bits per column, and TM-align: $>$ 0.5 TM-score) are shown by red-dashed lines.

**Foldseek**

| Model | Description | Homologous | Probability |
|---|---|---|---|
| AF-P03876-F1-v1 | PUTATIVE COX1/OXI3 INTRON 2 PROTEIN | ✓ | 1.0 |
| AF-P03875-F1-v1 | PUTATIVE COX1/OXI3 INTRON 1 PROTEIN | ✓ | 1.0 |
| AF-P0A3U1-F1-v1 | GROUP II INTRON-ENCODED PROTEIN LtrA | ✓ | 1.0 |
| AF-A0A0G2L2B6-F1-v1 | REVERSE TRANSCRIPTASE DOMAIN-CONTAINING PROTEIN | ✓ | 1.0 |
| AF-Q54ZK5-F1-v1 | REVERSE TRANSCRIPTASE DOMAIN-CONTAINING PROTEIN | ✓ | 1.0 |
| AF-Q55D49-F1-v1 | REVERSE TRANSCRIPTASE DOMAIN-CONTAINING PROTEIN | ✓ | 1.0 |
| AF-Q54E03-F1-v1 | REVERSE TRANSCRIPTASE DOMAIN-CONTAINING PROTEIN | ✓ | 1.0 |
| AF-P11369-F1-v1 | LINE-1 RETROTRANSPOSABLE ELEMENT ORF2 | ✓ | 1.0 |
| AF-P0A3U0-F1-v1 | GROUP II INTRON-ENCODED PROTEIN LtrA | ✓ | 1.0 |
| AF-B1N1A3-F1-v1 | PUTATIVE NICOTINE OXIDOREDUCTASE | ✓ | 1.0 |

**Foldseek-TM**

| Model | Description | Homologous | TM score |
|---|---|---|---|
| AF-A0A1D6L213-F1-v1 | RETROVIRUS-RELATED POL POLYPROTEIN LINE-1 | ✓ | 0.501 |
| AF-A0A1D8PFY7-F1-v1 | REVERSE TRANSCRIPTASE DOMAIN-CONTAINING PROTEIN | ✓ | 0.484 |
| AF-A0A0R0J4L8-F1-v1 | RNA-DEPENDENT RNA POLYMERASE | ✓ | 0.477 |
| AF-A0A1D6Q028-F1-v1 | RETROVIRUS-RELATED POL POLYPROTEIN LINE-1 | ✓ | 0.474 |
| AF-A0A1D6ERU1-F1-v1 | REVERSE TRANSCRIPTASE DOMAIN-CONTAINING PROTEIN | ✓ | 0.465 |
| AF-Q54LM7-F1-v1 | REVERSE TRANSCRIPTASE DOMAIN-CONTAINING PROTEIN | ✓ | 0.465 |
| AF-E7F608-F1-v1 | REVERSE TRANSCRIPTASE | ✓ | 0.465 |
| AF-Q2QZT5-F1-v1 | REVERSE TRANSCRIPTASE | ✓ | 0.459 |
| AF-P38478-F1-v1 | UNCHARACTERIZED MITOCHONDRIAL PROTEIN YMF40 | ✓ | 0.457 |
| AF-Q47688-F1-v1 | PROTEIN YkfC | ✓ | 0.456 |

**TMalign**

| Model | Description | Homologous | TM score |
|---|---|---|---|
| AF-A0A1D6L213-F1-v1 | RETROVIRUS-RELATED POL POLYPROTEIN LINE-1 | ✓ | 0.501 |
| AF-A0A1D8PFY7-F1-v1 | REVERSE TRANSCRIPTASE DOMAIN-CONTAINING PROTEIN | ✓ | 0.484 |
| AF-A0A0R0J4L8-F1-v1 | RNA-DEPENDENT RNA POLYMERASE | ✓ | 0.477 |
| AF-A0A1D6JF34-F1-v1 | RETROVIRUS-RELATED POL POLYPROTEIN LINE-1 | ✓ | 0.474 |
| AF-A0A1D6Q028-F1-v1 | RETROVIRUS-RELATED POL POLYPROTEIN LINE-1 | ✓ | 0.473 |
| AF-A0A1D6G0V4-F1-v1 | RETROVIRUS-RELATED POL POLYPROTEIN LINE-1 | ✓ | 0.473 |
| AF-A0A1D6H1V1-F1-v1 | RETROVIRUS-RELATED POL POLYPROTEIN LINE-1 | ✓ | 0.473 |
| AF-A0A1D6LAQ1-F1-v1 | RETROVIRUS-RELATED POL POLYPROTEIN LINE-1 | ✓ | 0.473 |
| AF-A0A1D6K060-F1-v1 | RETROVIRUS-RELATED POL POLYPROTEIN LINE-1 | ✓ | 0.472 |
| AF-A0A1D6ERU1-F1-v1 | REVERSE TRANSCRIPTASE DOMAIN-CONTAINING PROTEIN | ✓ | 0.466 |

**Dali**

| Model | Description | Homologous | Z-score |
|---|---|---|---|
| AF-P92543-F1 | UNCHARACTERIZED MITOCHONDRIAL PROTEIN ATMG01110 | ✓ | 11.7 |
| AF-Q3EC48-F1 | MITOVIRUS RNA-DEPENDENT RNA POLYMERASE | ✓ | 11.0 |
| AF-K7LVI8-F1 | UNCHARACTERIZED PROTEIN | ✓ | 10.3 |
| AF-K7LE20-F1 | UNCHARACTERIZED PROTEIN | ✓ | 10.2 |
| AF-P03876-F1 | PUTATIVE COX1/OXI3 INTRON 2 PROTEIN | ✓ | 10.2 |
| AF-Q62120-F1 | TYROSINE-PROTEIN KINASE JAK2 | ✓ | 9.9 |
| AF-K7K434-F1 | UNCHARACTERIZED PROTEIN | ✓ | 9.8 |
| AF-A0A1D6P530-F1 | PROTEIN KINASE DOMAIN CONTAINING PROTEIN | ✓ | 9.6 |
| AF-A0A1D6GQJ0-F1 | NON-SPECIFIC SERINE/THREONINE PROTEIN KINASE | ✓ | 9.6 |
| AF-Q54E07-F1 | REVERSE TRANSCRIPTASE DOMAIN-CONTAINING PROTEIN | ✓ | 9.6 |

**Supplementary Table** 3: **RdRp search** The search for homologous structures of RdRps (6M71_A) was done with Foldseek, Foldseek-TM module and TM-align against the AlphaFold/Proteome and AlphaFold/Swiss-Port database. Dali was only searched through the AlphaFold/Proteome database due its long runtime. The top 10 hits of the respective method are shown. We labeled if structures are homologous by manually checking the Uniprot website for RdRp/RT or Kinase domains.

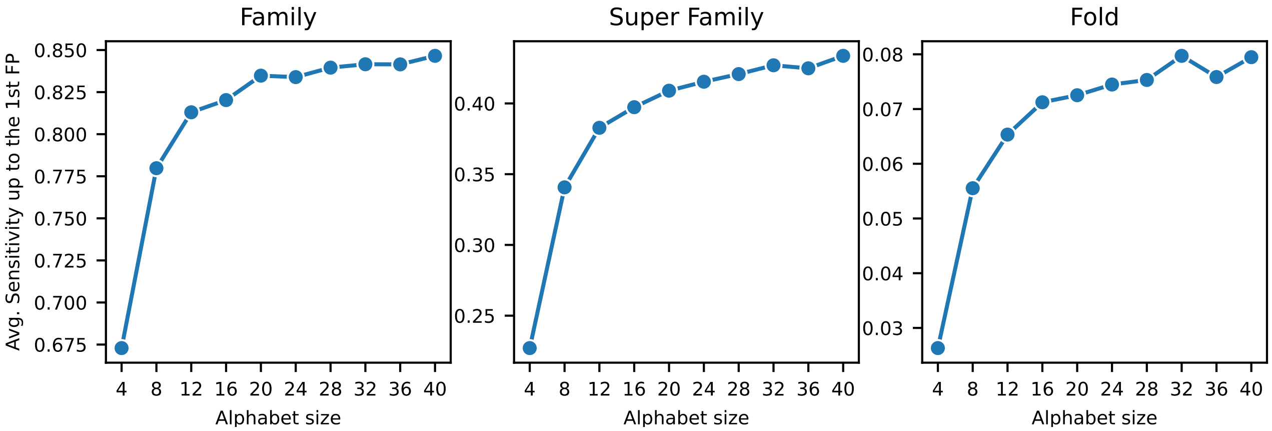


**Supplementary Figure** 13: **SCOPe benchmark sensitivity by 3Di alphabet size** We evaluated the effect of the alphabet size by training a new alphabet (VQ-VAE and substitution matrix) and testing it on the SCOPe benchmark for each size. The average sensitivity up to the 1st FP is the area under the ROC curve, as in the benchmark in **FIG. 2**.

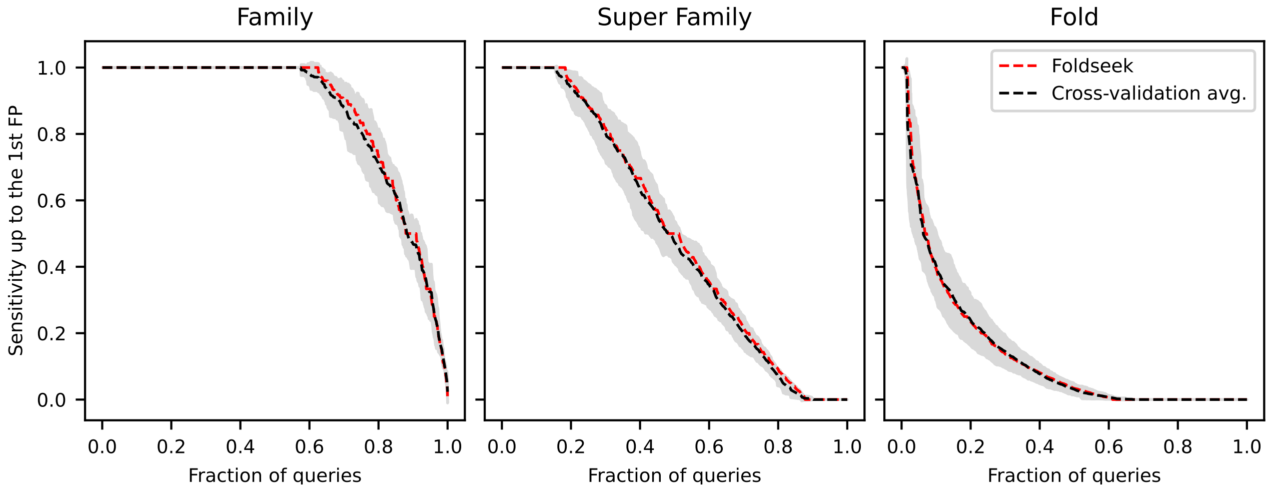


**Supplementary Figure** 14: **3Di alphabet: 4-fold cross-validation on SCOPe40** The 3Di alphabet was trained and benchmarked on the same dataset (SCOPe40). We therefore tested the generalization capabilities of the model with a 4-fold cross-validation, where the SCOPe40 dataset was split by fold into four parts (all domains of each fold are in the same split). Four new models were trained on three parts and benchmarked on the remaining part. The figure shows their mean sensitivity (black) and the standard deviation (gray area) in comparison to the final 3Di alphabet that is used in Foldseek (red), which was trained on the entire dataset.
